# Expressions for Bayesian confidence of drift diffusion observers in dynamic stimuli tasks

**DOI:** 10.1101/2020.02.25.965384

**Authors:** Joshua Calder-Travis, Rafal Bogacz, Nick Yeung

**Affiliations:** Department of Experimental Psychology, University of Oxford; MRC Brain Network Dynamics Unit, Nuffield Department of Clinical Neuroscience, University of Oxford

**Keywords:** Perceptual decisions, confidence, DDM, Bayesian, dynamic stimuli

## Abstract

We introduce a new approach to modelling decision confidence, with the aim of enabling computationally cheap predictions while taking into account, and thereby exploiting, trial-by-trial variability in dynamic stimuli. Using the framework of the drift diffusion model of decision making, along with time-dependent thresholds and the idea of a Bayesian confidence readout, we derive expressions for the probability distribution over confidence reports. In line with current models of confidence, the derivations allow for the accumulation of “pipeline” evidence that has been received but not processed by the time of response, the effect of drift rate variability, and metacognitive noise. The expressions are valid for stimuli that change over the course of a trial with normally-distributed fluctuations in the evidence they provide. A number of approximations are made to arrive at the final expressions, and we test all approximations via simulation. The derived expressions contain only a small number of standard functions, and require evaluating only once per trial, making trial-by-trial modelling of confidence data in dynamic stimuli tasks more feasible. We conclude by using the expressions to gain insight into the confidence of optimal observers, and empirically observed patterns.

## 1 Introduction

How humans and other animals make perceptual decisions is of fundamental interest. It is increasingly recognised that decision confidence, an estimate of the probability a decision was correct, is both theoretically important and used in a variety of ways to shape individual and group decision making (Bahrami et al., 2010; Desender, Boldt, Verguts & Donner, 2019; Desender, Boldt & Yeung, 2018; Drugowitsch, Mendonça, Mainen & Pouget, 2019; Sanders, Hangya & Kepecs, 2016; van den Berg, Zylberberg, Kiani, Shadlen & Wolpert, 2016). Confidence has also been linked to psychological disorder (Hauser, Allen, Rees & Dolan, 2017; Rouault, Seow, Gillan & Fleming, 2018). Reflecting the significance of confidence judgements, substantial efforts have been made to characterise their underlying computational mechanisms (e.g. Balsdon, Wyart & Mamassian, 2020; Desender, Donner & Verguts, 2020; Fleming & Daw, 2017; Geurts, Cooke, van Bergen & Jehee, 2022; Kiani, Corthell & Shadlen, 2014; Moran, Teodorescu & Usher, 2015; Pleskac & Busemeyer, 2010; Ratcliff & Starns, 2009; van den Berg, Anandalingam et al., 2016; Yu, Pleskac & Zeigenfuse, 2015; Zylberberg, Barttfeld & Sigman, 2012). The present work builds on these efforts to introduce a set of mathematical expressions for confidence using the drift diffusion model (also known as the diffusion decision model; DDM; Ratcliff and McKoon, 2008), coupled with a Bayesian readout for confidence (Kiani & Shadlen, 2009; Moreno-Bote, 2010; Sanders et al., 2016). The novelty of our contribution is our combination of three aims: To derive expressions for confidence within normatively prescribed frameworks for decision making and metacognitive evaluation; to go beyond simply fitting to aggregated confidence reports and instead develop methods capable of tractable fits to (and predictions about) individual confidence reports (Park, Lueckmann, von Kriegstein, Bitzer & Kiebel, 2016); and to provide a flexible modelling framework that can incorporate (and estimate the impact of) several factors recently shown or hypothesised to influence confidence reports. A key feature of our approach is that we derive expressions for the probability distribution over confidence reports (given decisions and response times) rather than focusing on first-order decisions and response-times themselves. Such expressions can be used as the basis for model fitting and parameter estimation, while avoiding the computational cost of making trial-by-trial predictions for decisions and response times (Ratcliff, 1980; Shan, Moreno-Bote & Drugowitsch, 2019; Smith, 2000; Smith & Ratcliff, 2021).

It is important to ask at the outset why we would want to derive explicit mathematical expressions that are computationally cheap to evaluate, when computational modelling can often be performed in other ways. This exercise has two main purposes: We aim for computationally cheap predictions to make trial-by-trial modelling of dynamic stimuli feasible, and we aim for explicit mathematical expressions to gain deeper insight into the nature of confidence in an important model. Explicit mathematical expressions provide immediate knowledge of the relationships between different parameters and variables, and how they combine to produce confidence. Indeed, we will see that we can go further and interpret such expressions to understand why different variables have the effect they do (Section 4). Regarding our other motivation, making trial-by-trial modelling feasible, such modelling allows us to capitalise on variability in stimuli, rather than ignoring it or treating it as noise (Park et al., 2016). When modelling on a trial-by-trial basis, a model that can capture the specific effects of each stimulus will outperform a model that can only capture general patterns in confidence across conditions (such as condition means or distributions of aggregated confidence reports). Hence, computationally cheap expressions may facilitate the development and testing of models for confidence that make increasingly precise predictions for behaviour.

The DDM is one of the most prominent models of two-alternative decisions, from a family of models in which observers receive noisy measurements of evidence for the two options (Bogacz, Brown, Moehlis, Holmes & Cohen, 2006; Green & Swets, 1966; Ratcliff & McKoon, 2008). In the DDM, observers track the difference in evidence measurements between the two options. That is, for each sample, observers subtract the measurement for option B from the measurement for option A, and add this to a running total (Ratcliff & McKoon, 2008). When the accumulator tracking this difference reaches a fixed threshold (positive or negative), a response is triggered. The DDM has successfully been used to model decisions in a wide range of tasks (Ratcliff, Smith, Brown & McKoon, 2016).

The DDM is also a normative model of decision making. By “normative” and “optimal” we refer to observers that, using the information assumed available to them, maximise reward rate (Rahnev & Denison, 2018). Under certain conditions, such as evidence measurements for the two options being equally reliable and signal strength known, the DDM is equivalent to tracking the posterior probability of each option until a fixed threshold on these probabilities is reached (Bitzer, Park, Blankenburg & Kiebel, 2014; Gold & Shadlen, 2007; Moran, 2015). In such a context, this policy maximises reward rate (Moran, 2015; Wald & Wolfowitz, 1948). When signal strength is unknown, a time-dependent threshold is required, but under standard assumptions it nevertheless remains optimal to track the difference between the two accumulators (Drugowitsch, Moreno-Bote, Churchland, Shadlen & Pouget, 2012; Moran, 2015; Tajima, Drugowitsch, Patel & Pouget, 2019). For an intuition of why this policy is optimal, consider a case in which the observer hasn’t made a decision after lengthy deliberation. The observer must be accumulating evidence very slowly, suggesting to them that signal strength is very low. If the observer thinks signal strength is very low, there is almost nothing to gain from collecting more evidence measurements, so they should lower their decision threshold and make an immediate decision (Malhotra, Leslie, Ludwig & Bogacz, 2017).

Although the DDM has optimal characteristics, and has been successfully applied to a wide range of decisions (Ratcliff et al., 2016), it is not clear the DDM provides an adequate account of confidence reports. Different ways of modelling confidence using the DDM have been proposed (Yeung & Summerfield, 2014). In one set of models, observers use some form of heuristic, based on variables which are directly accessible in the DDM, such as the state of the accumulator (Pleskac & Busemeyer, 2010), or the time taken to make a decision (Zylberberg et al., 2012). Another approach is to assume that observers map the state of the accumulator, and the time spent accumulating evidence, to the probability they are correct (Kiani et al., 2014; Kiani & Shadlen, 2009; Moreno-Bote, 2010). A Bayesian readout of this kind could be learned over time, through the association of accumulator state and time with success or failure (Kiani & Shadlen, 2009). Alternatively, a Bayesian readout could reflect a probabilistic inference made using knowledge of the statistical structure of the task. One detail to consider is that the confidence readout could be based on a separate evidence accumulator to the one used for the decision, or on multiple evidence accumulators (Balsdon et al., 2020; Fleming & Daw, 2017; Ganupuru, Goldring, Harun & Hanks, 2019; Jang, Wallsten & Huber, 2012; Ratcliff & Starns, 2009, 2013). However, here we make the simplest assumption that decisions and confidence are based on the same, single, normative, evidence accumulator (Moreno-Bote, 2010).

There are several techniques that can be used to calculate the probability, according to the DDM, of different responses and response times (Brown, Ratcliff & Smith, 2006; Chang & Cooper, 1970; Cox & Miller, 1965; Diederich & Busemeyer, 2003; Drugowitsch, 2016; Navarro & Fuss, 2009; Shinn, Lam & Murray, 2020; Smith, 2000; Tuerlinckx, 2004; Tuerlinckx, Maris, Ratcliff & De Boeck, 2001; Voss & Voss, 2008). Importantly, approaches have been developed that can handle dynamic stimuli (stimuli that change over the course of a trial) and time-dependent thresholds. One approach involves using finite difference methods to approximate the evolution of the probability distribution over accumulator state (which reflects the accumulated difference in evidence measurements; Chang and Cooper, 1970; Shinn et al., 2020; Voss and Voss, 2008; Zylberberg, Wolpert and Shadlen, 2018). Time and space are discretised and, working forward from the first time step, we solve a set of simultaneous equations at each time step to find the evolution of the probability distribution over accumulator state. If we are only interested in the probability distribution over response times and choices, we can use expressions described by Smith (2000). To evaluate these expressions we only need to discretise time, not space. Again working forward from the first time step, we can calculate the probability of deciding at each time step. Both approaches require that we discretise the time course of a trial into small time steps, and the number of computations required will scale with the number of time steps considered. Hence, in both approaches, we must perform a large number of computations (unless we use a task and stimulus in which modelling with large time steps is justified; Park et al., 2016).

This computational cost becomes important if we want to leverage the trial-by-trial variability inherent in dynamic stochastic stimuli. (Practical solutions for trial-by-trial modelling already exist for static stimuli under certain conditions; e.g. Wiecki, Sofer and Frank, 2013). Often the computation time needed for calculating predictions is reduced by making predictions on a condition-by-condition basis (Park et al., 2016; e.g. Kiani et al., 2014; Ratcliff and McKoon, 2008; van den Berg, Anandalingam et al., 2016; Zylberberg, Fetsch and Shadlen, 2016). We design the experiment so that there are a small number of different conditions (e.g. levels of stimulus contrast), then we treat all trials from a single condition as the same, and make predictions for behaviour in each condition, rather than for each stimulus individually. This approach does not capitalise on the trial-by-trial variability of the dynamic stimuli that are often used (for model-free analyses that do capitalise on trial-by-trial variability see Charles and Yeung, 2019; Kiani, Hanks and Shadlen, 2008; Zylberberg et al., 2012). A method that reduced the computational cost of making trial-by-trial predictions for dynamic stimuli could allow us to perform model fitting that capitalises on rather than discards the rich data produced from such stimuli.

Another approach is to derive explicit mathematical expressions for model predictions, which could dramatically reduce computation time. Moreno-Bote (2010) derived expressions for confidence that take into account time-dependent thresholds, and that could be extended to account for dynamic stimuli of the kind we consider below. However, these derivations use two assumptions about the computation of confidence which are not in line with recent findings. First, Moreno-Bote (2010) made the intuitive assumption that decisions and confidence are based on the same information. However, as sensory and motor processing takes time, there will be stimulus information in these processing “pipelines” that does not contribute to the initial decision, but that nevertheless informs subsequent confidence judgements (Charles & Yeung, 2019; Moran et al., 2015; Ratcliff & McKoon, 2008; Resulaj, Kiani, Wolpert & Shadlen, 2009; van den Berg, Anandalingam et al., 2016). Moreover, information in the processing pipeline is affected by trial-to-trial fluctuations in signal strength, and this may have important effects on confidence (Pleskac & Busemeyer, 2010). Second, there is now substantial evidence that the process that “reads out” confidence into a behavioural report is corrupted by “metacognitive noise” (Bang, Shekhar & Rahnev, 2019; De Martino, Fleming, Garrett & Dolan, 2013; Maniscalco & Lau, 2012, 2016; van den Berg, Yoo & Ma, 2017). This additional noise in the readout may reflect imperfections in the transfer of information between decision and metacognitive processes (Bang et al., 2019; De Martino et al., 2013), or may be because confidence reports themselves are sensitive to factors outside of the first-order decision process (Maniscalco, McCurdy, Odegaard & Lau, 2017; Rahnev, Koizumi, McCurdy, D’Esposito & Lau, 2015; Shekhar & Rahnev, 2020). We aim for mathematical expressions that take these important features into account.

A key idea that affects the scope of our derivations is that, whereas it may be very difficult or impossible to derive simple expressions for decisions and response times, it may be much simpler to find expressions for confidence (given a specific decision and response time). Prior to a decision, even if changes to the state of the accumulator are normally distributed over small intervals of time, the probability distribution over the state of the accumulator will not be normal. This is because, having reached a time *t* without a response, we know that the accumulator is not beyond either threshold, nor has it been at any point up to *t* (otherwise the observer would already have made a decision; Moreno-Bote, 2010). This constraint results in non-normal probability distributions over accumulator state (Fig. 1A), with associated mathematical expressions that either feature infinite summations, or may even be intractable (Ratcliff, 1980). In contrast, it is much simpler to express confidence directly, in terms of the evolving state of the accumulator following a decision. Ratcliff (1980) previously reported that we can expect a normal distribution over the state of the evidence accumulator when decision thresholds are absent, even if evidence signal strength varies within a trial (as would be expected from dynamic stimuli). We build on this work by considering confidence in the related situation of evidence accumulation following a decision threshold crossing. This situation turns out to be more complex, nevertheless – following a decision – decision boundaries are no longer relevant, hence normally distributed changes in the state of the accumulator lead to a normal distribution over this state (Fig. 1B). Thus, our contribution is not to provide new expressions for the probability distribution over response times and decisions in diffusion models. We aim to bypass the complexities associated with response times and decisions, and instead focus on confidence.

**Figure 1:**
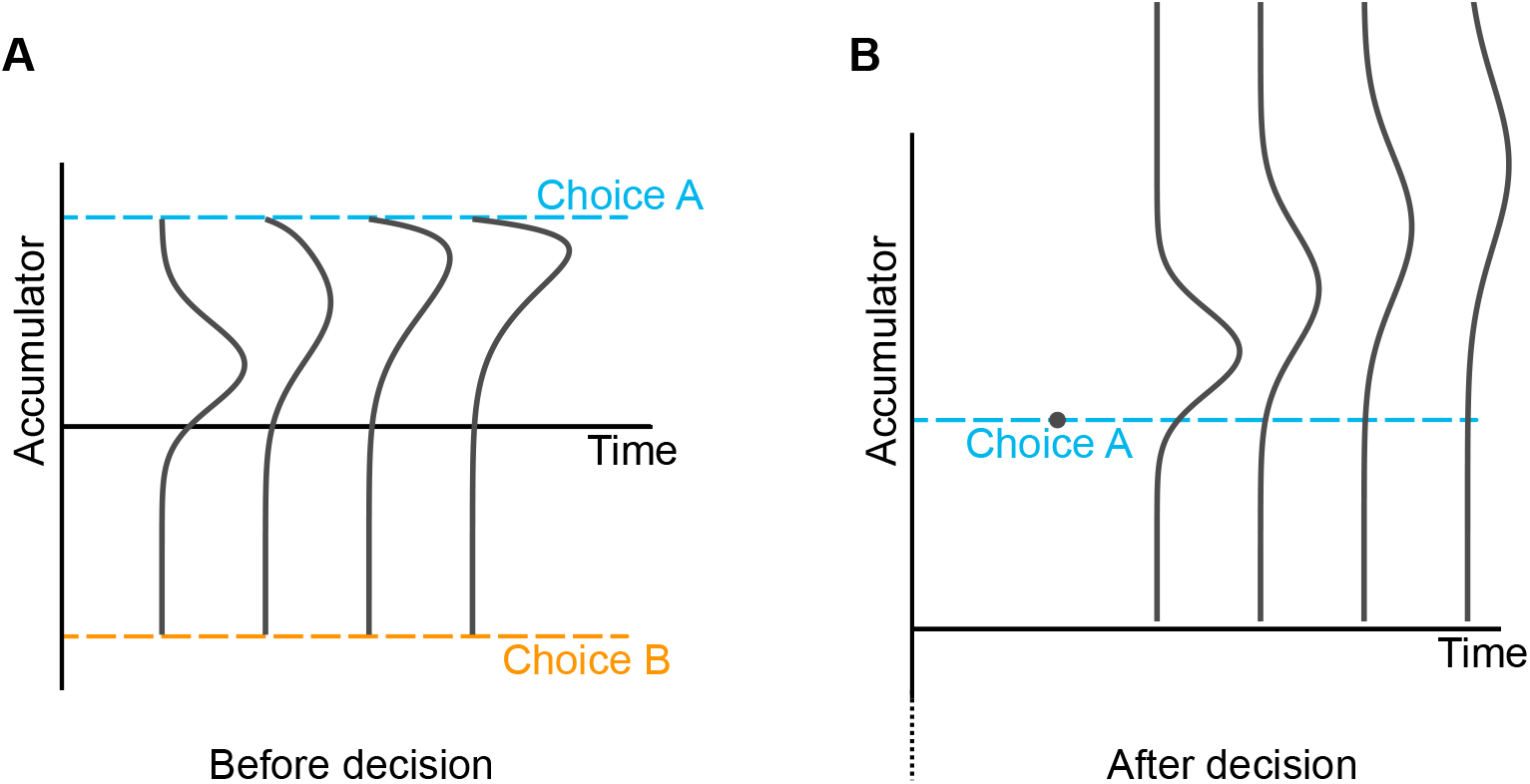
Probability distributions over accumulator state before and after a decision. Even if increments in the accumulator are normally distributed, prior to a decision the probability distribution over the state of the accumulator quickly becomes non-normal. This is because, if we get to time *t* without a decision, we know the accumulator has not been beyond either decision threshold prior to *t*. Following a decision (the time of which is represented by the dot in the right panel), normally distributed increments in the accumulator lead to a normal distribution over accumulator state, because there are no longer decision thresholds.

Using this strategy we derive approximate expressions for the probability, according to the DDM, of different confidence reports, given the response and response time on a trial. As discussed, the framework of the DDM with possibly time-dependent thresholds, includes (under standard assumptions) the optimal decision policy, whether or not signal strength is known by the observer. Confidence is allowed to be a noisy readout of the probability of being correct, and we account for the effects of pipeline evidence, and variability in signal-to-noise ratio. The derived expressions only need evaluating once per trial, instead of at each very small time step within each trial, and allow for dynamic stimuli of a certain form, thereby making trial-by-trial modelling of such stimuli feasible. As discussed, making trial-by-trial modelling feasible is one of our main aims. Once we have derived mathematical expressions for confidence, we will additionally be able to use them to gain deeper insight into the nature of confidence within the framework of the normative DDM.

## 2 Model

### Overview

Our aim is to derive expressions for the probability distribution over confidence, given the response and response time on a trial. In order to derive these predictions we must first specify a model for decisions and confidence, and the context in which that model is to be applied. The equations in the following subsections formalise a generic two-alternative decision making task with time-varying evidence, and specify a model that conforms to the well-established ideas of the DDM (Ratcliff & McKoon, 2008; Ratcliff et al., 2016). This DDM-based model features some natural extensions – inspired by previous work – to deal with the possibility of time-varying evidence (Drugowitsch et al., 2012; Ratcliff, 1980; Smith, 2000). Finally, a confidence readout is specified, which is based on the common idea that observers read out the probability they are correct (Kiani et al., 2014; Moran, 2015; Pew, 1969; Sanders et al., 2016).

We consider a situation in which observers must make a choice between two alternatives. The presented stimulus provides two evidence signals, one for each option, and evidence provided by the stimulus can vary over time (Fig. 2; Bogacz et al., 2006; Moreno-Bote, 2010). For example, the observer might be presented with two clouds of dots, with the number of dots in each cloud constantly changing. Their task might be to determine which of the two clouds contains the most dots on average (Charles & Yeung, 2019). Here the dots in the two clouds would correspond to the two streams of evidence. We assume that the observer only receives noisy measurements of the presented evidence (Green & Swets, 1966; Ratcliff & McKoon, 2008). In our model, consistent with the DDM, the observer takes the difference between the noisy measurements of evidence for the two options, and accumulates this difference (Fig. 2).

**Figure 2:**
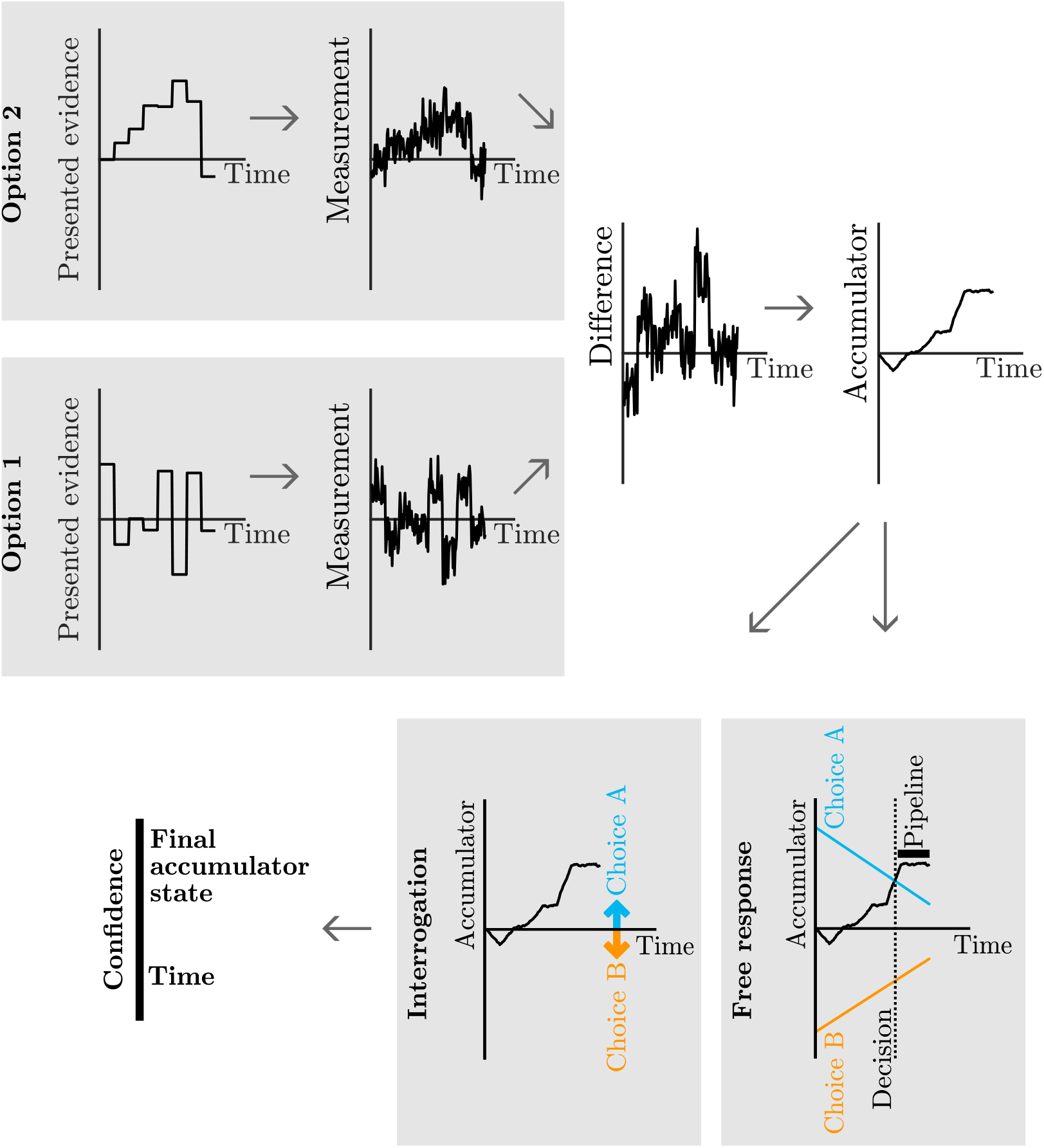
The model of confidence and decisions. Time varying evidence is presented for the two response options. The observer only receives noisy measurements of the evidence presented. The observer takes the difference between the evidence measurements and accumulates this difference. In the interrogation condition, the duration of the stimulus is set by the researcher. The observer accumulates noisy measurements until the stimulus ends and all evidence is processed. Then the observer simply picks the option which is favoured by the accumulated measurements. In the free response condition, the observer uses decision thresholds, one for each option, which trigger a response. Following a decision, evidence measurements in the processing pipeline are accumulated, and confidence is informed by the accumulator state at the time of threshold crossing, plus changes to the accumulator caused by evidence measurements from the pipeline.

Typically, following stimulus onset, a participant can respond whenever they wish. Some instruction or incentive may be given to respond in a certain way, such as fast and accurately, but beyond this the participant is free to set the time of response (e.g. Ratcliff & McKoon, 2008). The stimulus continues to be presented until a response is made. We call this condition “free response” (but it is also referred to as the “information controlled” condition elsewhere; Bogacz et al., 2006; McMillen and Holmes, 2006; Ratcliff, 1980). Following the DDM, we assume the observer sets two thresholds on the accumulator state, one for each choice (Bogacz et al., 2006; Ratcliff & McKoon, 2008). When the accumulator reaches one of these thresholds, the corresponding response is triggered (Fig. 2). As discussed in Section 1, measurements corresponding to evidence that has recently been presented will still be in sensory and motor processing pipelines at the time of response, and hence will not contribute to a decision (Resulaj et al., 2009). These measurements will be processed immediately following a response, and will be used to inform confidence (van den Berg, Anandalingam et al., 2016).

We also consider the “interrogation” condition (McMillen & Holmes, 2006), where the observer must respond at a time controlled by the researcher. (This and closely related conditions are also referred to as “time controlled” and “response signal” conditions; Dosher, 1976; Ratcliff, 1980, 2006; Schouten and Bekker, 1967; Usher and McClelland, 2001.) In this case the stimulus is presented for a finite amount of time. Before the stimulus clears the participant cannot respond. Once the stimulus clears, the observer uses the final state of the accumulator (which reflects all evidence presented in the stimulus) to determine their response (Fig. 2; Bogacz et al., 2006). Although we will focus on this normative decision making strategy (Bogacz et al., 2006), we note that an alternative non-normative modelling choice would be to also include the idea of decision thresholds in the interrogation condition, assuming that if a decision threshold is reached people stop accumulating evidence and withhold their response until the appropriate time (Balsdon et al., 2020; Ratcliff, 2006). There is evidence that careful selection of when response times are enforced in the interrogation condition can minimise this possibility (Rosenbaum, Glickman, Fleming & Usher, 2022). In both the free response and interrogation conditions, the observer uses a Bayesian readout of confidence which depends on the final state of the accumulator once all evidence has been processed, and the time spent accumulating evidence (Kiani et al., 2014; Moreno-Bote, 2010).

The following subsections are organised as follows. First, we describe the observer’s task mathematically, and the beliefs held by the observer. Second, we find the rule a Bayesian observer would use to map evidence measurements to a decision and confidence. Third, we test the effect of the beliefs we ascribe to the observer. Fourth, we describe the noisy “read out” process which determines confidence (Fleming & Daw, 2017). In section 3 we turn to the main aim of the paper, which is to use the drift diffusion framework to derive simple expressions for the probability distribution over confidence reports, given a response and response time. A summary of symbols used in the derivations can be found in Table 1. We use the convention that log refers to the natural logarithm.

**Table 1:**
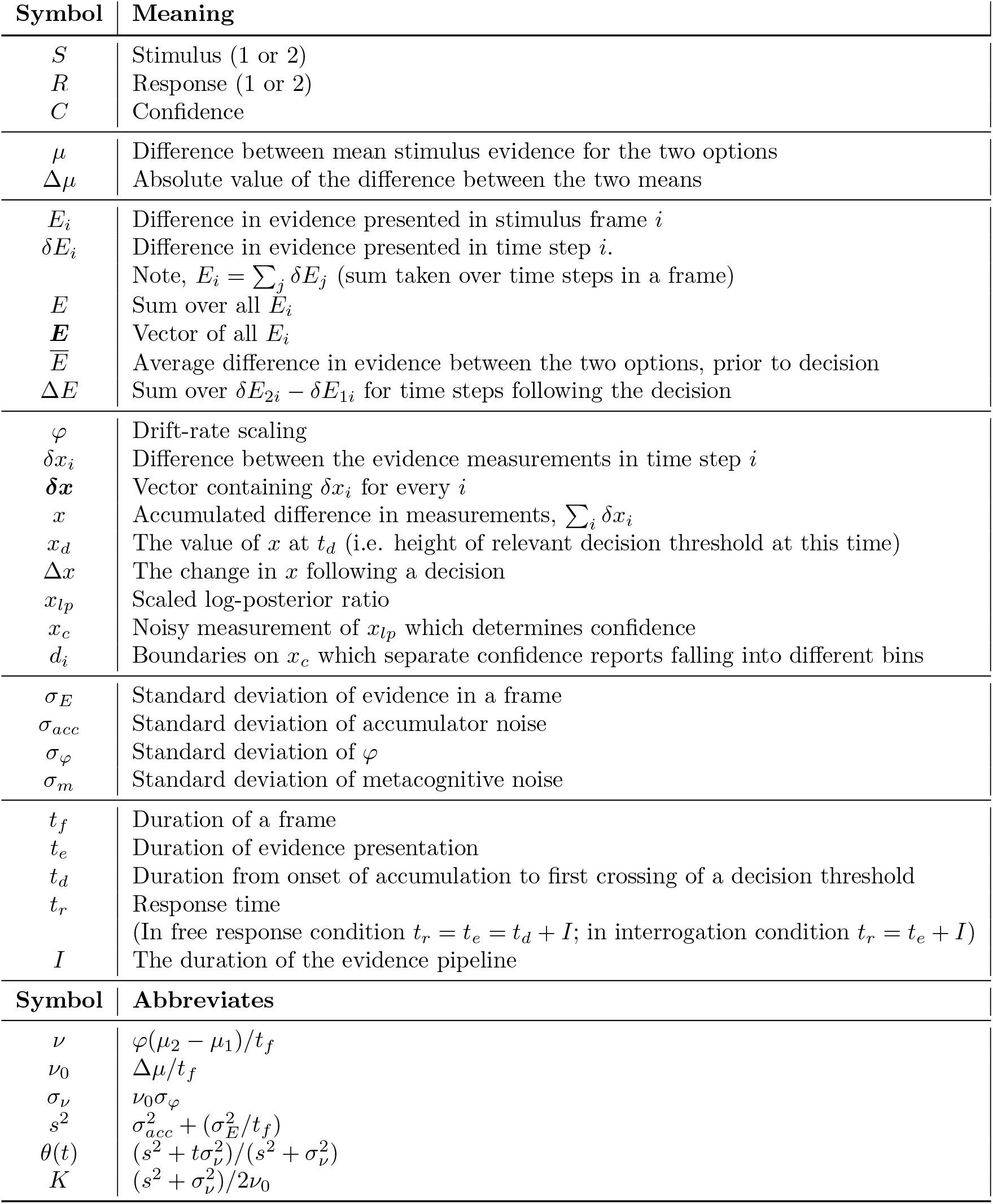
Symbols used in the derivations, along with key abbreviations.

### Task and observer

The observer’s task is to determine the correct response, by inferring which evidence stream is drawn from the distribution with the greater mean (Bogacz et al., 2006; Moreno-Bote, 2010). As we are considering a DDM observer who tracks the difference in the evidence measurements for the two alternatives, we only need to consider the difference in evidence provided by the two evidence signals from the stimulus.

We consider the case where the two options are equally probable. Denote the mean evidence for the two options, *μ*_1_ and *μ*_2_, and the difference between these means, *μ* = *μ*_2_ − *μ*_1_. We consider a situation in which the absolute value of the difference between the two means is a fixed value. However, we incorporate variability in signal strength below (see discussion of variability in drift-rate scaling, *φ*). Let Δ*μ* denote a fixed positive value which determines the absolute value of the difference between the two means. This setup gives us,

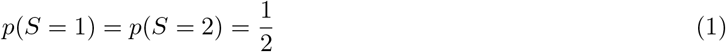

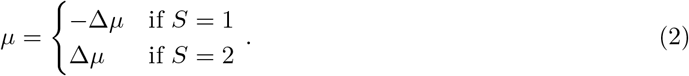

*S* denotes the stimulus (1 or 2) with the greater mean evidence, and hence the correct answer.

Denote the total time during which the stimulus is presented *t_e_*, and the time of the response relative to the beginning of the trial *t_r_*. In the free response condition *t_e_* = *t_r_* because a response triggers the end of the stimulus. For the free response condition we have to consider the effects of the decision thresholds (discussed above). Denote the time spent accumulating measurements prior to the first crossing of a decision threshold, *t_d_*. A threshold crossing triggers a response. However, because of delays in sensory and motor processing, there is a lag between the point of internal commitment to a decision and the point at which that decision is externally registered via an overt movement, and even at the point of the overt response some recently received sensory evidence will still be being processed (Luce, 1986; Resulaj et al., 2009). Hence, information presented in the stimulus immediately prior to the response will not contribute to the decision. Denote the duration of the stimulus immediately prior to the response that does not contribute to the decision, because corresponding measurements are still in processing pipelines, *I*. The time spent accumulating evidence until the first decision threshold crossing and the duration of the processing pipeline, will together equal the time taken to respond, *t_r_* = *t_d_* + *I*.

In the interrogation condition the observer uses all information presented in the stimulus – by accumulating evidence for a duration equal to the duration of evidence presentation – to determine both their response and confidence (Bogacz et al., 2006). Once all evidence has been accumulated, after *t_e_*, a response is then triggered. We make the natural assumption that sensory and motor processing delays are of the same duration in both the interrogation and free response conditions, although in the interrogation condition no further information is gathered during this time because the stimulus is no longer being presented. In this condition, response time is therefore given by *t_r_* = *t_e_* + *I*.

We consider here the general case of a stimulus that provides time-varying evidence (Fig. 2). Our derivations also apply to time-constant evidence as a special case of time-varying. We consider evidence that is piecewise-constant within short stimulus “frames” of duration *t_f_*. We use *E_i_* to denote the evidence presented for option 2 minus the evidence presented for option 1 (i.e., the difference in evidence) during frame *i*. If the difference in evidence presented in each frame is normally distributed around the underlying mean then we have,

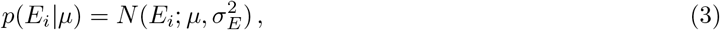

where 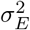 is the variance over the presented difference-in-evidence. Note that *μ* does not correspond to anything the observer directly observes. *μ* is the underlying mean difference-in-evidence presented for the two options, and is constant throughout a trial. The actual presented evidence, *E_i_*, is what varies over the course of a trial, and is drawn from a distribution centred on the underlying means.

Although observers could in principle leverage knowledge that evidence is constant throughout each frame, we assume that observers ignore this additional structure. This will be a valid approximation when evidence frames used in an experiment are sufficiently short. In this case, the approximation will lead to little discrepancy between the observer’s estimate of the probability of being correct, and the true probability. We test this claim below, once we have derived an expression for the probability of being correct (Fig. 3, and equation 20). Consider a discretisation of time into very short time steps (much shorter than the duration of a frame) of duration *δt*. Instead of accounting for the constant evidence over the course of a frame, we assume that observers treat the fraction of evidence received in a short time step, *j*, as generated according to,

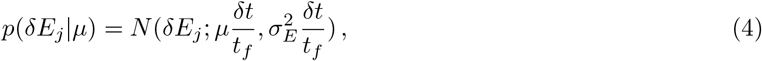

where, *δE_j_* indicates the difference-in-evidence presented in time step *j* only, not over the course of a frame (*E_i_* = ∑*_j_ δE_j_* where the summation is taken over all the time steps in a stimulus frame). Importantly, the observer incorrectly assumes that the *δE_j_* at different time steps, *j*, are independent (i.e., the observer ignores the fact that evidence is constant through all time steps in a single frame).

**Figure 3:**
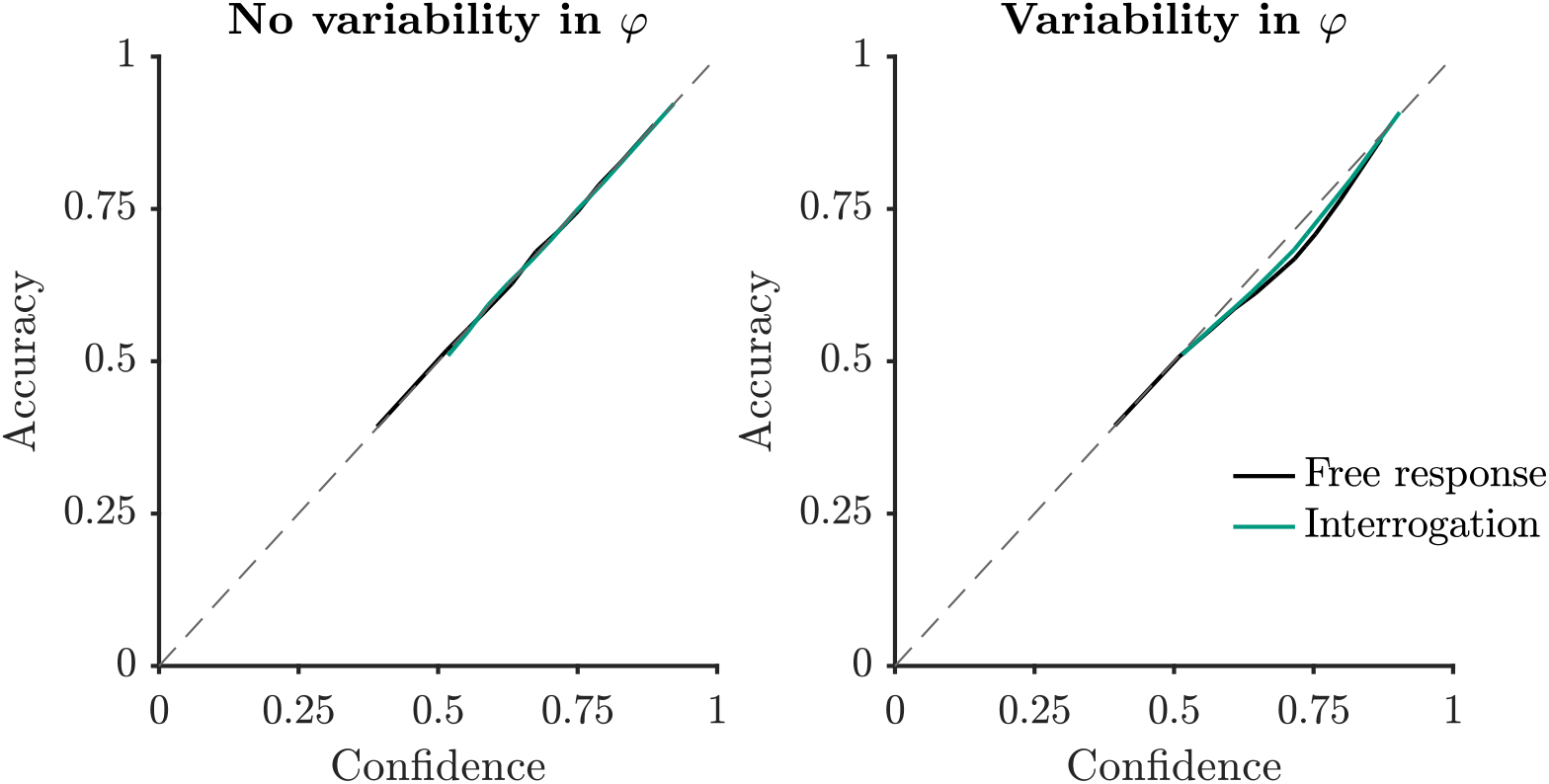
Accuracy as a function of confidence, where confidence is calculated using the observer’s generative model of their evidence measurements. This model only approximates the true generative model. When there is variability in drift-rate scaling, the approximations cause the observer’s confidence to show some deviations from calibration. Simulation and plotting details in Appendix D.

As in the DDM, the presented evidence drives an internal evidence accumulation that is subject to normally distributed noise (Ratcliff & McKoon, 2008). We also take into account the fact that the stimulus varies over time: Each stimulus frame drives the evidence accumulation for a duration equal to the duration that frame is presented for. Over a small time step, *j*, the incremental change in the state of the accumulator tracking the difference in evidence measurements, denoted *δx_j_*, can be described by,

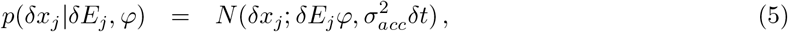

where 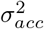 is the variance of noise in the accumulation (Drugowitsch et al., 2012). *δE_j_* is determined by the stimulus frame currently being processed. *φ* is a random variable which accounts for variability in drift rate. “Drift rate” refers to the rate at which evidence presented in the stimulus drives the accumulation of evidence measurements (Ratcliff & McKoon, 2008). Where a stimulus, and hence drift rate, is constant over the course of a trial, drift rate variability is trial-to-trial variability in this rate (Ratcliff, 1978; Ratcliff & McKoon, 2008; Ratcliff & Smith, 2004; Voskuilen, Ratcliff & Smith, 2016). Here, where stimulus evidence, and hence drift rate, varies over the course of each trial, we operationalise drift rate variability as a multiplicative factor that determines how well stimulus information is processed. To distinguish this operationalisation from the usual operationalisation, we sometimes refer to the multiplicative factor *φ* as the “drift-rate scaling”, because we set the mean value of this variable to be one regardless of the strength of the presented evidence. When the drift-rate scaling is high, the signal extracted from the stimulus is greater, and evidence is accumulated rapidly. Noise in the accumulation is unaffected, hence, a higher drift-rate scaling also leads to a higher signal-to-noise ratio.

It is usually assumed that drift rate variability follows a normal distribution (Ratcliff, 1978; Ratcliff & McKoon, 2008). We make the same assumption here,

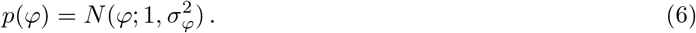

We assume that observers correctly model the effect of internal noise (accumulator noise, and drift-rate scaling variability). We return to the plausibility of this assumption in the discussion. Hence observers infer that increments in the accumulator are related to the underlying signal as follows,

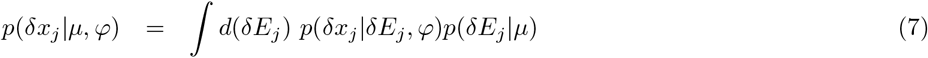

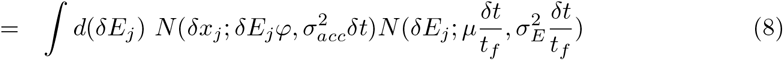

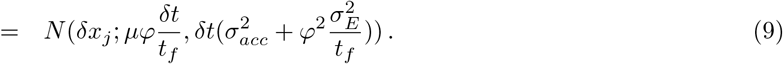

To reach the final line we rearranged and applied the result in Appendix A. In words, this equation states that evidence measurements have a mean that matches the underlying stimulus signal multiplied by the drift-rate scaling, but are variable due to both internal accumulator noise and the fact that evidence in the stimulus is itself variable. It is interesting to note that the drift-rate scaling, *φ*, not only affects the mean increment, but also the variability of increments, through the term 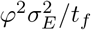. This effect arises because high drift-rate scaling increases the effect of the stimulus on the accumulation, increasing the effects of both the stimulus mean and variability in the stimulus. We assume that participants ignore this difference between stimulus variability and internal variability (which is not affected by drift-rate scaling). Hence we assume that observers believe evidence measurements are corrupted by the same amount of variability regardless of the level of drift-rate scaling:

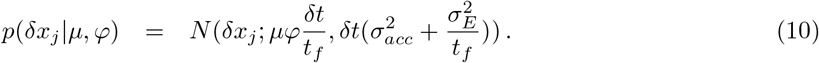

We test the effect of this assumption on the calibration of confidence below (Fig. 3).

To simplify these expressions we change variables to 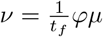, and 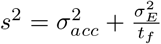. This gives,

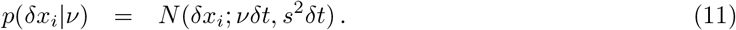

We can compute the probability distribution over *ν* given *S* using (2) and (6), giving,

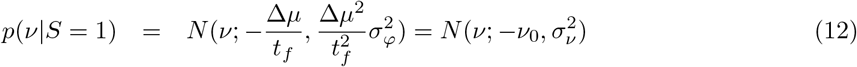

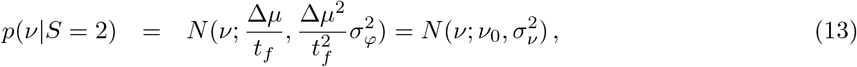

where 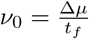, and 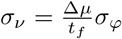.

In summary, observers are aiming to infer *S*, which affects *ν* via,

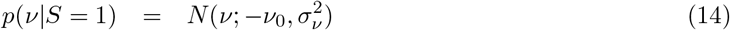

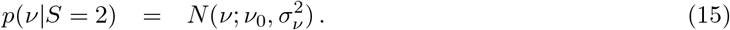

They receive two independent streams of evidence. They look at the difference in evidence in these streams at each time point, a quantity which they assume is related to *ν* via,

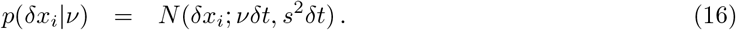

### The Bayesian observer

Having specified the properties of the task and the beliefs of the observer, we can infer the rule used by a Bayesian observer to translate all evidence measurements gathered so far into a decision and confidence. We denote the vector of all evidence measurements, *δx*_1_, *δx*_2_, *…, δx_N_* as *δx*. Inferring the posterior probability over *S*, given *δx*, in the environment described by (14), (15), and (16), is an inference problem that has been considered before. Moran (2015), using a result from Drugowitsch et al. (2012), derived the posterior probability over the two options *S* = 1 and *S* = 2. We also provide a derivation in Appendix B, and simply state the result here. Using Bayes rule with all evidence samples the log-posterior ratio is given by (Appendix B; Moran, 2015),

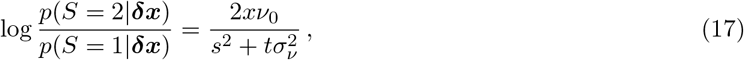

where 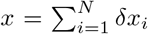, and *t* is the time spent accumulating evidence. We will often work with a scaled version of the log-posterior ratio,

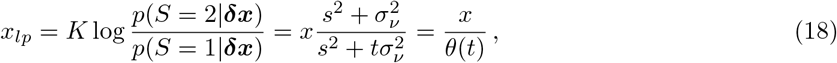

where *x_lp_* denotes the scaled version of the log-posterior ratio. *θ*() provides an abbreviation for the purpose of making future equations less cluttered. *K* is the scaling constant and is equal to,

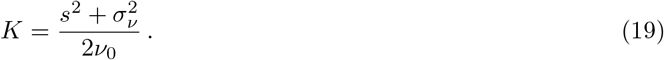

Note that the log-posterior ratio without any scaling is given by *x_lp_/K*.

A Bayesian observer would report whichever option is more likely. Hence, they will report *S* = 1 when *x_lp_/K* < 0, which is the same as *x_lp_* < 0, or *x* < 0. Denote this report *R* = 1, and a report for *S* = 2 as *R* = 2. In the interrogation condition, the observer simply has to wait for the stimulus to end at *t_e_*. Once the observer has processed and accumulated all evidence measurements, the observer can respond according to the sign of the final accumulator state, *x* (Fig. 2). In the free response condition, the observer uses a decision threshold for triggering a response (Bogacz et al., 2006; Ratcliff & McKoon, 2008). This threshold describes an absolute value of the accumulator, |*x*|, which triggers a response when reached. We allow the threshold to vary with time (Drugowitsch et al., 2012). As discussed, at the time of response, measurements corresponding to recently presented evidence will still be in the processing pipeline (Resulaj et al., 2009). The response will be based on *x* at the time of the decision, *t_d_*, while confidence will incorporate additional pipeline evidence measurements (Fig. 2). Processing will continue until measurements from the full duration of stimulus presentation, *t_e_*, have been processed.

The observer can use the (scaled) log-posterior ratio in the computation of confidence, as it is monotonically related to the probability they are correct through,

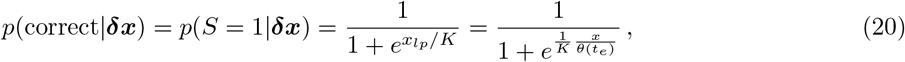

if the observer reports *R* = 1. (This expression can be found by rearranging *p*(*S* = 1*|**δx***)+*p*(*S* = 2*|**δx***) = 1.) A similar expression holds for *R* = 2, the only difference being that *x* is replaced with −*x*, and *x_lp_* with −*x_lp_*.

### Testing observer model approximations

Above, we assumed that observers approximate the true generative model for the evidence measurements. Specifically we assumed, (a) observers ignore the fact that evidence is constant within each frame, and (b) observers ignore the increased effect of variability in the stimulus, when the drift-rate scaling is high, and the decreased effect when the drift-rate scaling is low. While the assumptions may be plausible, it is also plausible that, if given feedback, observers could learn to closely approximate the mapping from *x* and *t_e_* to probability correct (Kiani & Shadlen, 2009). To adequately capture this situation we need to ensure that, under our assumptions about the observer, confidence remains closely related to performance.

To test the two approximations, we simulated the diffusion process using small time steps. At each time step the accumulator increment, *δx_i_*, was determined by drawing a value consistent with (5), until a decision threshold was crossed (Tuerlinckx et al., 2001). Accumulation continued until all evidence measurements had been processed, at which point a simulated confidence was determined in accordance with (20). Full details of the simulation can be found in Appendix D.

Fig. 3 shows accuracy against confidence for two cases, one in which there is no variability in drift-rate scaling, *φ*, and one with variability. Full details of the process used to plot the data are also provided in Appendix D. Only one approximation is relevant when there is no variability in drift-rate scaling, approximation (a). This approximation appears to have little effect on the mapping between accuracy and confidence. An additional approximation is relevant when there is variability in drift-rate scaling, assumption (b). We can see that this causes a moderate mismatch between accuracy and confidence (Fig. 3).

It is interesting to note that the latter approximation, that observers ignore the effect of drift rate on stimulus variability, is implicitly present in much previous work. This is because dynamic stimuli are often used, but stimulus variability is not treated separately from accumulator variability in the computational models applied (e.g. Kiani et al., 2014; Ratcliff and McKoon, 2008; van den Berg, Anandalingam et al., 2016, although see Zylberberg et al., 2016). Fig. 3 suggests this approximation does not cause large issues.

### Confidence reporting model

Having described the task, observer beliefs, tested those beliefs, and found the confidence of a Bayesian observer, we turn to the question of exactly what quantity confidence reports reflect. While the (scaled) log-posterior ratio, *x_lp_* is monotonically related to the probability of being correct, we do not assume that confidence is a direct readout. Instead, we allow the possibility that metacognitive noise corrupts this estimate (De Martino et al., 2013; Maniscalco & Lau, 2012, 2016), and hence that confidence is based on a noisy representation of *x_lp_*, denoted *x_c_*,

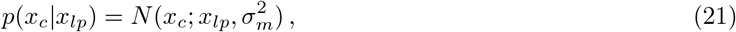

where *σ_m_* is the standard deviation of metacognitive noise. (Note that this noisy readout of the scaled log-posterior ratio, is equivalent to a scaled version of a noisy readout of the unscaled log-posterior ratio; Appendix C.)

We would also like to minimise the number of assumptions we make about how *x_c_* is transformed into a confidence report. There is evidence that different people use confidence scales in different ways (Ais, Zylberberg, Barttfeld & Sigman, 2016; Festinger, 1943; Navajas et al., 2017). To make minimal assumptions about how people view and treat confidence scales, we treat confidence reports, *C*, as ordinal data only (Aitchison, Bang, Bahrami & Latham, 2015). Confidence reports on a continuous scale can be analysed by binning them first.

If people report greater confidence when *x_c_* favours their decision to a greater extent, then all confidence reports falling into a higher confidence bin will have come from further along the *x_c_* scale (in the direction that favours the choice made). Using *d_i_* we denote the boundary on *x_c_* which separates the confidence reports which fall into confidence category *C* = *i −* 1 from *C* = *i*, when the observer reports *R* = 2. When *R* = 1, the boundary applies to −*x_c_*, or equivalently, a boundary of −*d_i_* is applied to *x_c_*.

## 3 Results

We now have a complete description of the model, and everything we need to derive the probability distribution over confidence reports in both the interrogation and free response conditions. We would like to find the probability distribution over confidence, given the evidence presented, ***E***, the response, *R*, and in the free response condition, the amount of time the observer monitors the stimulus before making a response, *t_r_*. (***E*** is a vector containing every *E_i_*.) A key variable is the observer’s (scaled) log-posterior ratio after they have seen all evidence, *x_lp_*. Our general strategy will be to find a probability distribution over this variable. From this distribution we will be able to infer a distribution over the noisy readout of the (scaled) log-posterior ratio, *x_c_*. As described in the previous section, on a trial with a response *R* = 2, if *x_c_* falls between *d_i_* and *d_i_*_+1_ the observer reports confidence *C* = *i*. If *R* = 1, the boundaries are −*d_i_*_+1_ and −*d_i_*. The probability of a confidence report *C* = *i* will be given by the probability that *x_c_* falls between the corresponding boundaries (Fig. 4).

**Figure 4:**
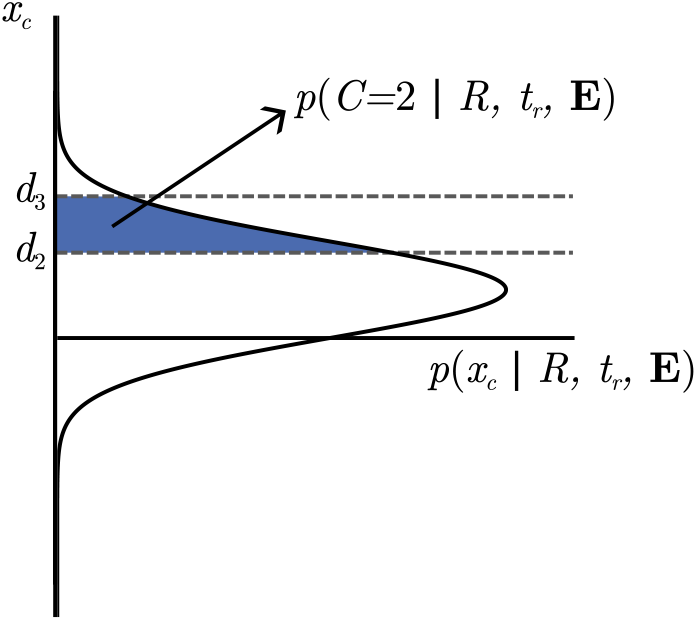
Probability of a confidence report, based on a distribution over *x_c_*. *x_c_* is a noisy representation of the (scaled) log-posterior ratio. We make no specific assumptions about how observers use confidence scales apart from assuming that, if the observer reports higher confidence *C*, then the underlying variable *x_c_* has a greater absolute value, in the direction corresponding to the response made. We use *d_i_* to denote the boundaries between values of *x_c_* that lead to different confidence reports. If we know the distribution over *x_c_*, then the probability of a specific confidence report can be found by integrating *x_c_* or −*x_c_* between the corresponding boundaries. Whether we integrate over *x_c_* or −*x_c_* depends on the response made.

Throughout we keep the dependence of the predictions on model parameters implicit: The probability distribution over confidence reports depends not just on the evidence presented, the response, and the time spent monitoring the stimulus, but also the parameters of the model. For the sake of readability this dependence is kept implicit in conditional probabilities in the derivations. (E.g. we write *p*(*C|R, t_r_, **E***) instead of *p*(*C R, t_r_, **E***, Ξ), where Ξ represents the set of parameters.) However the parameters on which the predictions depend are of course of great practical importance. It will be these parameters that we adjust as we fit the model to data, and by constraining particular parameters to certain values we will be able to create different variants of the model for comparison. Parameters for fitting to data are listed in Table 2. Decision threshold is listed but this is not in itself a parameter. We will see that the modeller has freedom over what shape decision threshold to use, and how to parameterise this function.

**Table 2:** Model parameters for fitting to data. The predictions for confidence depend on these parameters, but this dependence is kept implicit throughout for readability (i.e., a symbol for the set of parameters is not explicitly included when writing out conditional probabilities). The tables states “Parameter / feature” because *f* (*t_d_*) is not a parameter. *f* (*t_d_*) describes how the shape of the decision threshold changes over time. The modeller can parameterise this function as they wish. For example, they could use a flat threshold and simply fit threshold height, or they could use a complicated curved threshold with several parameters.

| Parameter / feature |  | Free response | Interrogation |
| --- | --- | --- | --- |
| Accumulator noise | $\sigma_{acc}$ | ✓ | ✓ |
| Drift-rate variability | $\sigma_{\varphi}$ | ✓ | ✓ |
| Metacognitive noise | $\sigma_m$ | ✓ | ✓ |
| Confidence boundaries | $d_i$ (for all $i$) | ✓ | ✓ |
| Duration of evidence pipeline | $I$ | ✓ | - |
| Decision threshold shape | $f(t_d)$ | ✓ | - |

### Interrogation condition

In a trial from the interrogation condition, the stimulus is presented for some amount of time, *t_e_*. The observer can only respond after the end of the stimulus. We aim to find the probability of confidence reports, given the response and evidence presented. Assuming the response occurs at some fixed amount of time following *t_e_*, the response time provides us with no information. This is because *t_e_* is set by the researcher, and hence unaffected by processes internal to the observer. A summary of the generative model for interrogation condition confidence reports, from the perspective of the researcher, is shown in Fig. 5.

**Figure 5:**
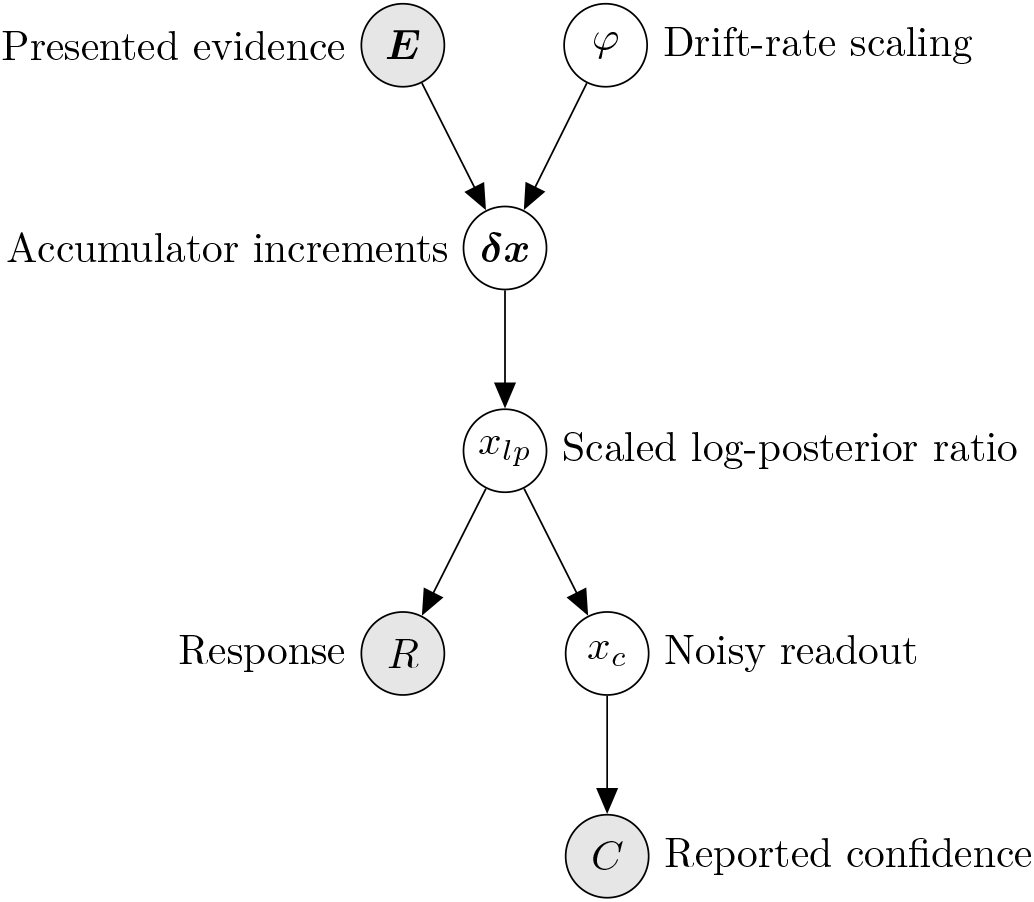
Representation of the generative model for confidence reports in the interrogation condition, from the perspective of the researcher. We want to infer the probability of reported confidence, using our knowledge of the evidence presented and the response given.

We start by integrating *x_c_* over the region which leads to a confidence report *C* = *i*. When the response is *R* = 2, this is the region between *d_i_* and *d_i_*_+1_ (Fig. 4). The case where *R* = 1 is identical except the limits become −*d_i_*_+1_ and −*d_i_*. Also marginalising over, *x_lp_* and using Bayes rule,

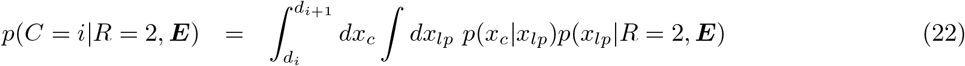

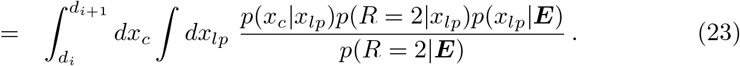

We want to find,

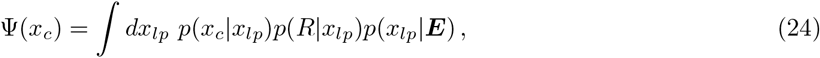

(where Ψ(*x_c_*) is an abbreviation), and then use,

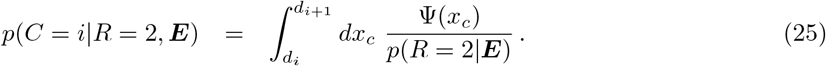

For now just consider *p*(*x_lp_|**E***). Let’s marginalise over *φ*,

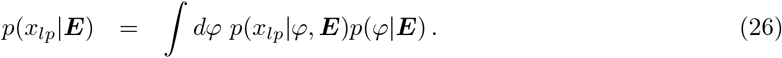

*E* and *φ* are independent of each other (when not conditioned on other variables; see Fig. 5). Hence,

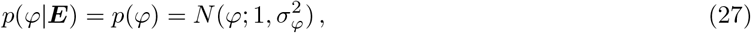

where we have used (6). Recall from (5) that,

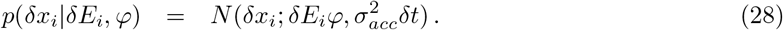

Using standard formulae for the sum of normally distributed random variables, we have that *x* = ∑*_i_ δx_i_* will be given by,

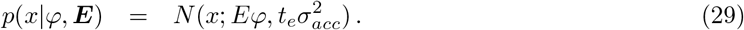

where *E* = ∑*_i_ δE_i_* (sum taken over all time steps in all relevant frames), and *t_e_* = ∑*_i_ δt*. Using (17) we have,

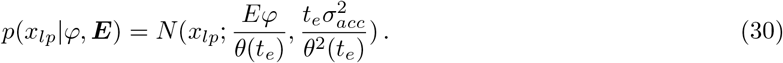

Substituting these results into (26) gives us,

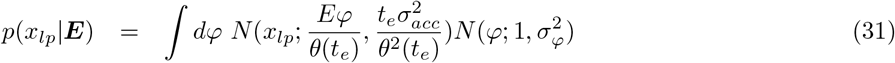

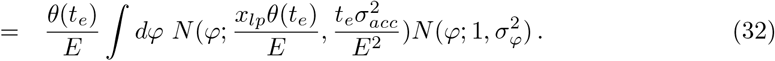

We can apply the result in Appendix A. Simplifying, we find,

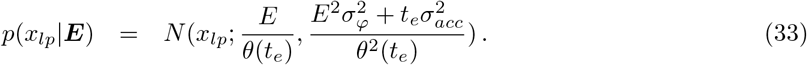

Returning to (24), we now have,

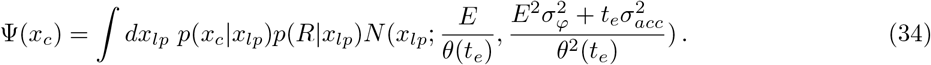

In the case in which we have no metacognitive noise, *x_c_* = *x_lp_*. We can express this using the Dirac delta function as *p*(*x_c_|x_lp_*) = *δ*(*x_c_ − x_lp_*). Then we have,

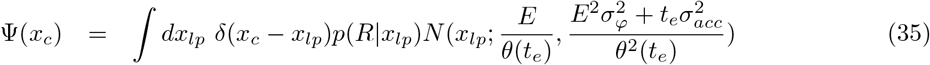

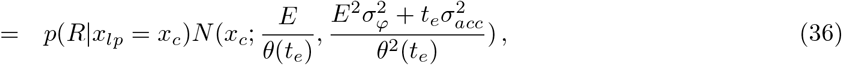

Consider the term *p*(*R|x_lp_* = *x_c_*), which describes the probability of a response given a final (scaled) log-posterior *x_lp_*. The Bayesian observer’s decision rule is deterministic, and was described in section 2. If *x_lp_* < 0 the observer reports that *R* = 1, and reports *R* = 2 if *x_lp_ >* 0. In all cases, the observer makes the response that is most likely to be correct. Hence, (scaled or not) the log-posterior ratio at the time of the decision always favours the response made, and the probability of *x_lp_* and *R* being inconsistent is zero. Due to the absence of metacognitive noise, *x_lp_* = *x_c_*. Hence, *p*(*R|x_lp_* = *x_c_*) will also be one when *x_c_* and *R* are consistent, and zero when *x_c_* is inconsistent with *R*, i.e., when *x_c_* suggests a different response should have been made. Because *x_c_* and *R* are always consistent, the observer will never report a confidence of less than 50%.

In the case with no metacognitive noise we have, using (25) and our expression for Ψ(*x_c_*),

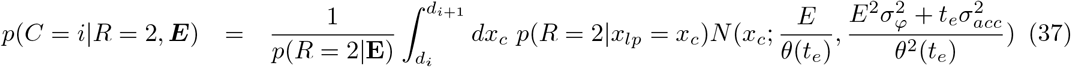

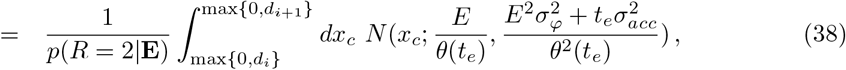

where we have used that *p*(*R* = 2*|x_lp_* = *x_c_*) is zero anywhere *x_lp_* is inconsistent with the response *R* = 2 (i.e. *x_lp_* < 0), and one elsewhere. In the case where *R* = 1 an identical expression holds, except the limits become min{0, −*d_i_*_+1_} and min{0, −*d_i_*}, and we divide by *p*(*R* = 1*|***E**) not *p*(*R* = 2*|***E**).

We can compute the probability of the response, *p*(*R|***E**), using,

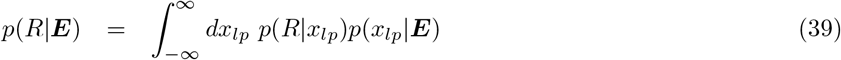

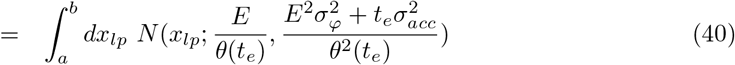

Where *a* = −*∞, b* = 0 for *R* = 1 or *a* = 0, *b* = *∞* for *R* = 2. These limits again come from using that *p*(*R|x_lp_*) = 1 when *R* and *x_lp_* are consistent, and *p*(*R|x_lp_*) = 0 otherwise.

Let’s now consider what happens in the presence of metacognitive noise, when *x_c_* no longer equals *x_lp_*, but is a noisy version of it as in (21). Returning to (34), in this case Ψ(*x_c_*) becomes,

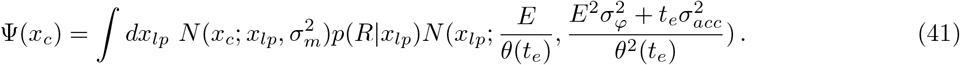

Because *p*(*R|x_lp_*) is one when *R* is consistent with the (scaled) log-posterior ratio, and zero otherwise, the product of this term and the normal distribution over *x_lp_* is a normal distribution truncated to the region where *R* and *x_lp_* are consistent. Note that the product of these two terms is not a probability distribution over *x_lp_* because it is not normalised. However, we can write this product as a scaled truncated normal distribution. Using *J* (*x_lp_*) as an abbreviation,

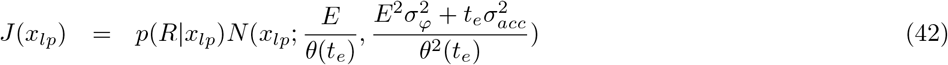

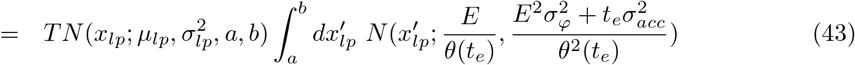

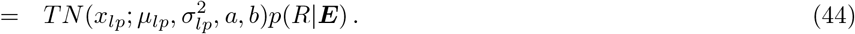

We have used (40) to get the final line. As before *a* = −∞, *b* = 0 or *a* = 0, *b* = ∞ depending on the response made, and *TN* indicates a truncated normal distribution, truncated at *a* and *b*. The first and second parameters, *μ_lp_* and 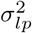, are the mean and variance of the distribution prior to truncation, and therefore are,

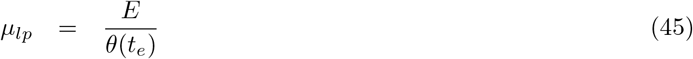

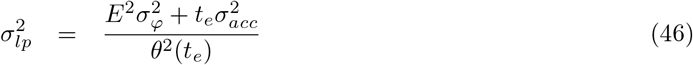

Returning to (41) we now have,

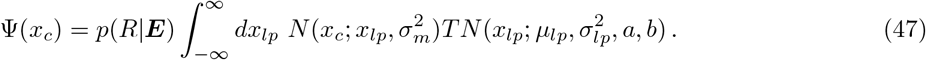

Consider a change of variables *y* = *x_c_ − x_lp_*,

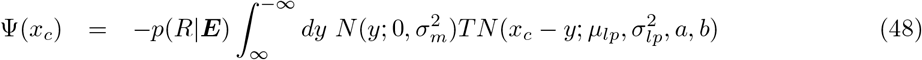

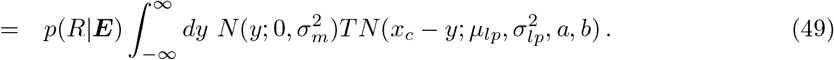

This expression is the convolution of a normal distribution and a truncated normal distribution. We can perform the convolution by writing the distributions out in full, collecting all terms containing *y* into a single exponential, and integrating (S. Turban, personal communication, December, 2019):

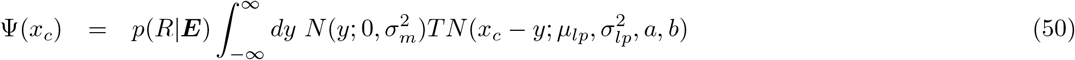

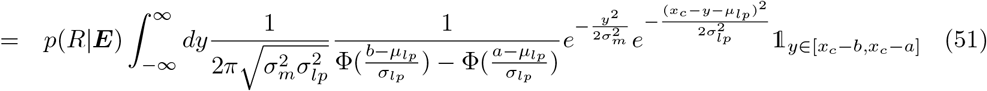

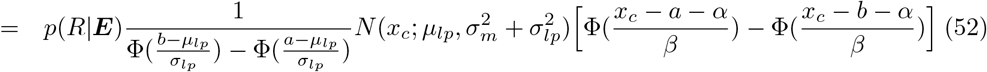

where Φ() indicates the standard cumulative normal distribution, 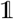 is an indicator function (which is one when its argument is true, and zero otherwise), 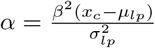 and 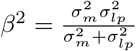.

In our case, *a* and *b* take very specific values. Let,

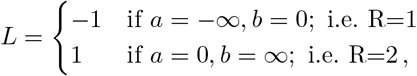

then we can write,

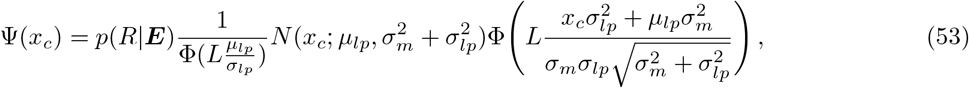

and using (25) our final result for confidence is,

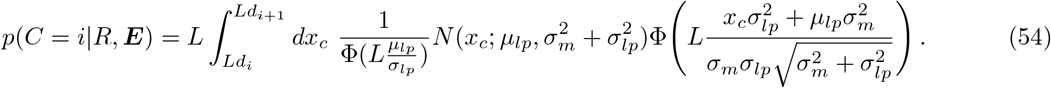

We have also used *L* in the above expression to account for the different limits of integration when *R* = 1 and *R* = 2.

We can solve the integral in (54) via numerical integration. It is also possible to rearrange the expression to a form that is faster to evaluate numerically. We rearrange the expression in Appendix E, and state the result here. The expression in (54) becomes,

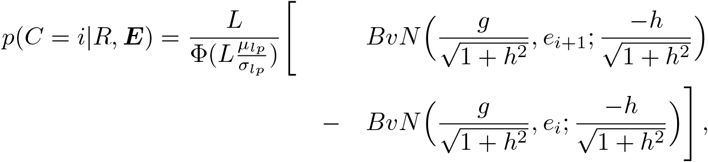

where,

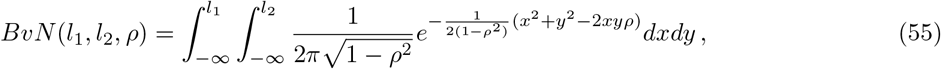

is the bivariate cumulative normal distribution, corresponding to a distribution with mean and covariance,

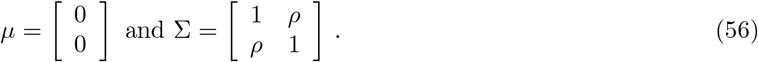

Additionally, *g*, *h*, and *e_i_* denote 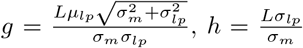 and 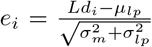. This integral can be numerically evaluated using standard functions.

### Free response condition

In the free response condition we again want to find the probability distribution over confidence reports, but now have an additional piece of information to incorporate into our predictions, the time of the response *t_r_*. Response time will be determined by the evolution of the accumulator, and specifically, by the first time the accumulator reaches a decision threshold (Fig. 6).

**Figure 6:**
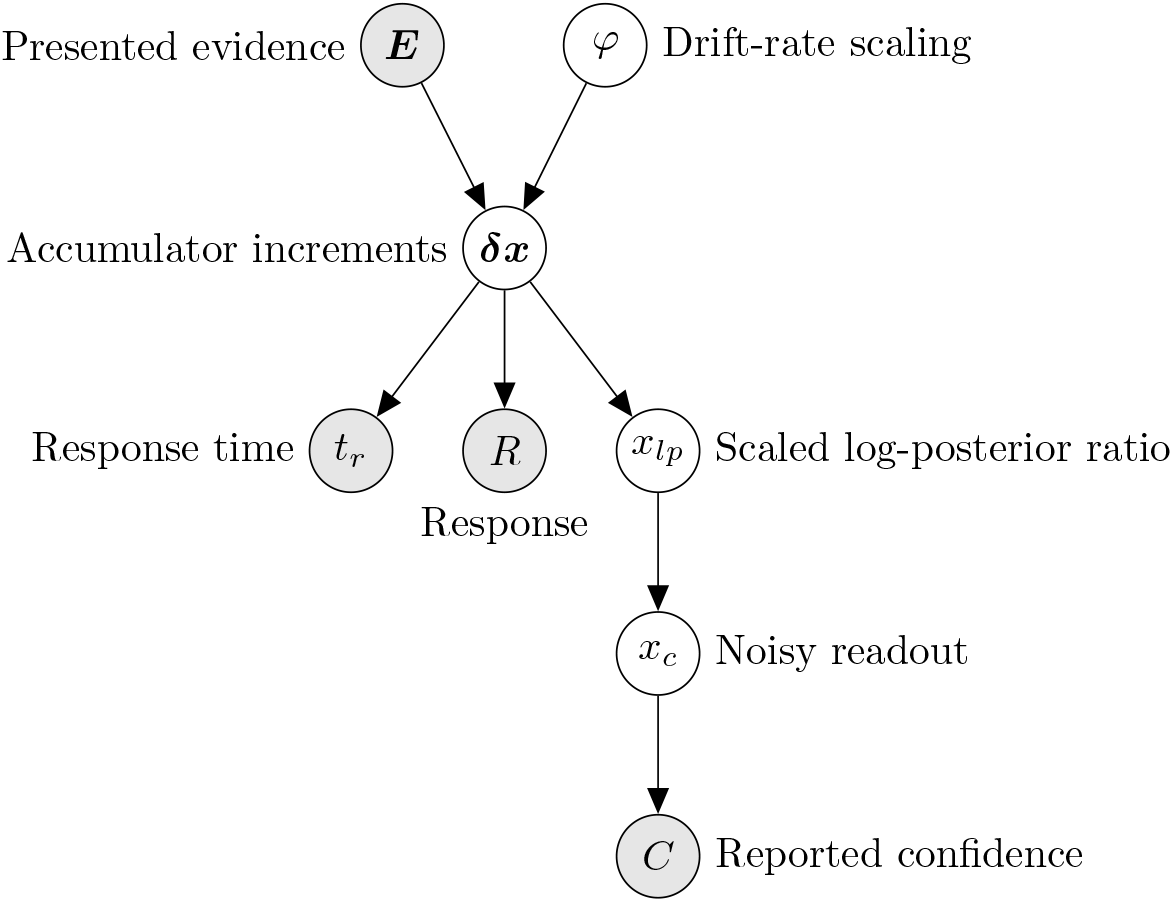
Representation of the generative model for confidence reports in the free response condition, from the perspective of the researcher. We want to infer the probability of reported confidence, using our knowledge of the evidence presented, the response given, and the response time.

Integrating *x_c_* over the region that leads to a confidence report *C* = *i* after *R* = 2, and marginalising over *x_lp_*,

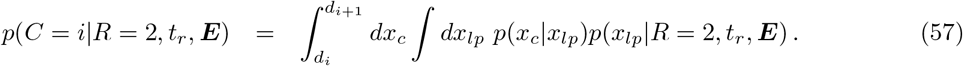

(As before, for *R* = 1 we use integration limits −*d*_*i*+1_ and −*d*_*i*_ instead.) The second distribution in this expression can be obtained by marginalising over *φ*,

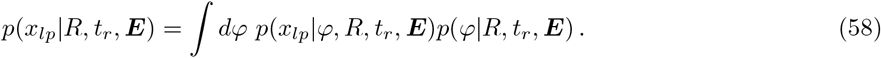

The final term is a distribution over *φ*, the drift-rate scaling. Observers often need to calculate the distribution over drift rate, given the evidence received (Drugowitsch et al., 2012; Moran, 2015; Moreno-Bote, 2010). As researchers, we can infer the value of *φ* in a similar way.

Evidence presented immediately prior to a response but after the decision point does not contribute to the response itself, as it is still being processed (Resulaj et al., 2009). If, for an observer, the interval of stimulus in this processing pipeline is *I*, we can infer that the amount of time they spent accumulating evidence prior to a decision, *t_d_*, is *t_d_* = *t_e_* − *I* = *t_r_* − *I*. Additionally, if an observer uses a decision threshold, *f* (*t*), to trigger responses *R* = 2, and −*f* (*t*) to trigger responses *R* = 1, then we know that at the time they made their decision *t_d_*,

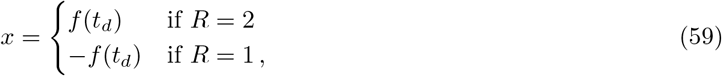

because the time they made their decision was the time the accumulator hit the decision threshold (Moreno-Bote, 2010). Denote the value of *x* at *t_d_* by *x_d_*. Hence for known (or hypothesised) values of *I* and *f* (*t*), we can infer *t_d_* and *x_d_* from *R* and *t_r_*. Note that in this case we can also infer *R* and *t_r_* from *t_d_* and *x_d_*. Hence, these quantities are interchangeable in

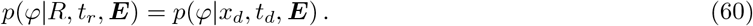

The probability distribution over *φ* will depend on the entire stream of evidence up to the time of the decision (i.e., all elements of ***E*** that correspond to evidence received before a decision). This is because measurements of all evidence prior to a decision, in conjunction with *φ*, determine changes in the accumulator, which in turn determine the time of the response, and response itself (see Fig. 6). However, for the purpose of inferring *φ*, we approximate the evidence stream by its average prior to the time of the decision, 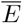,

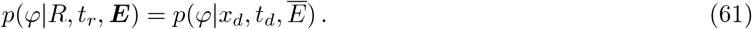

Precisely, 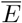 is given by 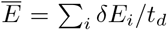, where the summation is taken over all time steps prior to the decision. We test this approximation, in conjunction with other approximations, once the derivation is complete. Using Bayes rule,

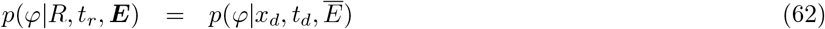

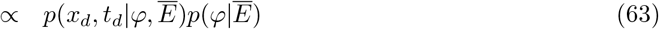

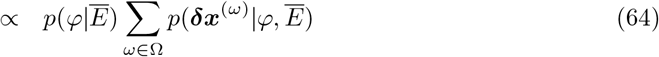

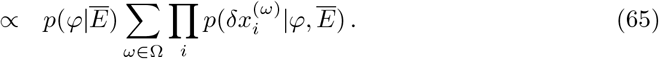

In this equation, Ω is the set of all paths that do not cross the decision threshold prior to *t_d_*, and arrive at the threshold at *t_d_* (Moreno-Bote, 2010). ***δx***^(*ω*)^ is the ***δx*** vector for the particular path *ω*.

We approximate 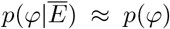. *φ* is independent of ***E***, but could depend on 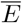, because the average evidence may be related to the time of the response, which *φ* also affects (Fig. 6). Again, this approximation will be tested once we have derived predictions for confidence. The approximation gives,

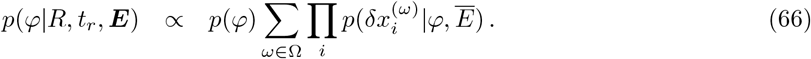

Returning to (5) but now using our approximation replacing the full evidence stream with the average evidence, we have that *δx_i_* is given by

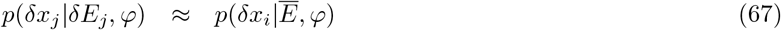

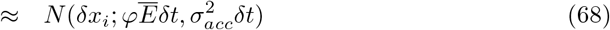

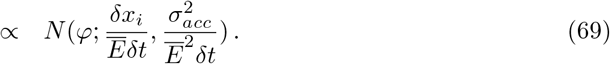

The proportionality is with respect to *φ*.

Using standard results for the product of normal distributions we find

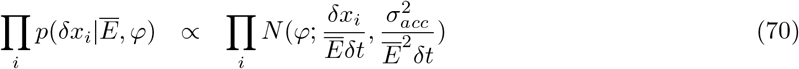

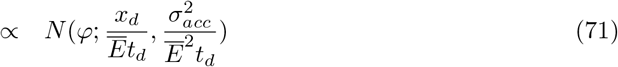

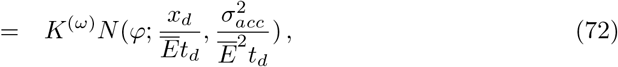

where we have used *t_d_* = ∑*δt*, *x_d_* = ∑*δx_i_*, and *K*^(*ω*)^ is the constant of proportionality (see Drugowitsch et al., 2012 for closely related calculations). This constant will be different for different paths, so we explicitly indicate the path, *ω*. Hence, we have in (66)

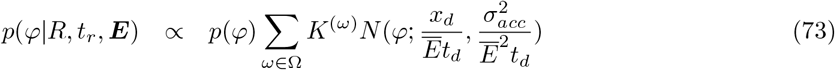

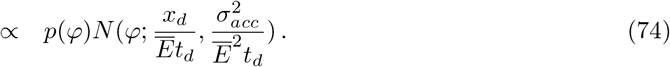

Recalling (6), *φ* is distributed as,

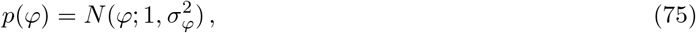

giving

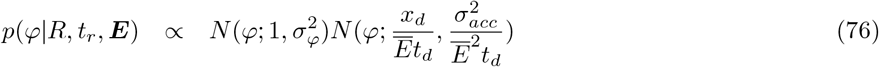

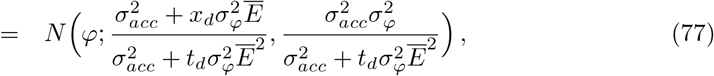

where we have used standard formulae for the product of normal distributions.

Returning to (58), we see that in addition to an expression for *p*(*φ|R, t_r_, **E***), we need an expression for *p*(*x_lp_ φ, R, t_r_, **E***). As discussed, from the response and response time, we can infer the time spent accumulating evidence prior to the decision, and the state of the accumulator at the time a decision was made (Fig. 2). Following a decision, evidence accumulation continues in the same manner as in the interrogation condition (Pleskac & Busemeyer, 2010), until all evidence measurements have been processed. Therefore, the expression in (29) is valid for predicting the accumulation between the time of the decision and the end of stimulus processing (*t_d_, t_e_*). Denote the accumulation in this time Δ*x*, and the sum over evidence presented in this interval Δ*E* = ∑*_i_ δE_i_* (the summation is taken over all *i* which correspond to time steps following a decision). Using (29),

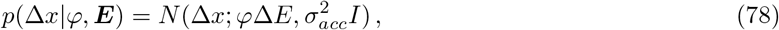

where *I* denotes the duration of the pipeline, *t_e_ − t_d_*. Using our knowledge of the location of *x* at *t_d_*, denoted *x_d_*, and that the final state of *x* is given by *x_d_* +Δ*x*, the distribution over the final state is given by,

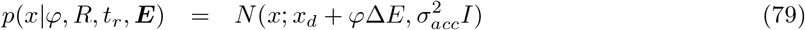

Using (17) we have,

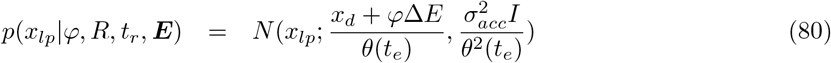

We have now derived expressions for both the distributions in (58). Returning to this equation we have,

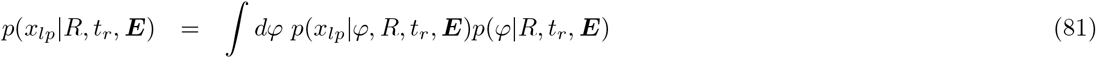

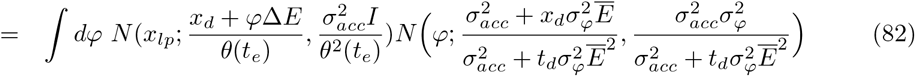

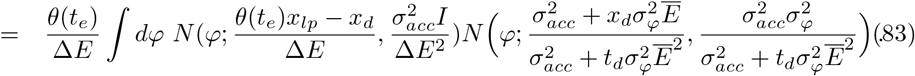

Applying the result in Appendix A and rearranging we find,

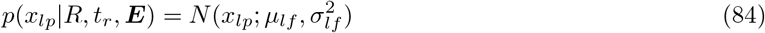

Where,

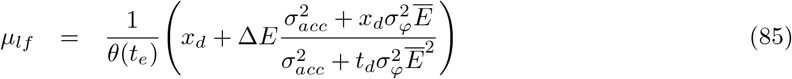

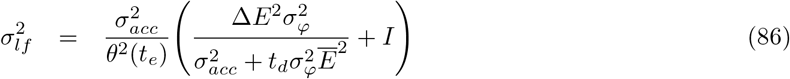

Using this result in (57), and allowing for normally distributed metacognitive noise as in (21),

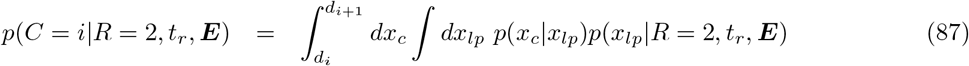

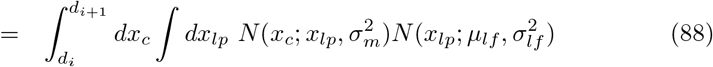

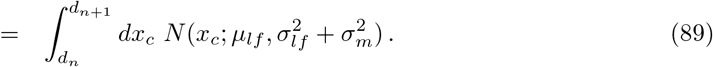

This expression applies for *R* = 2. For the case of *R* = 1 the limits change to −*d_i_*_+1_ and −*d_i_*.

Thus, we have derived expressions for the probability of confidence reports falling into each ordinal bin, given the trial-by-trial response, response time and stimulus. We have produced such expressions for both interrogation (54) and free response (89) decision tasks. By allowing for the possibility of various additional features in the underlying decision and confidence model, the derivations build on previous work, as set out in further detail in the discussion below. Equations (54) and (89) provide approximate expressions for confidence that only need evaluating once per trial, with the aim of supporting feasible trial-by-trial modelling even with time-varying stimuli.

### Testing the approximations

We made several approximations in the derivations above, so it is important to check that our predictions for confidence closely match confidence reports, when these are simulated. We simulated the diffusion process using small time steps, and produced confidence reports in accordance with the model (see (20), (21) and Fig. 2; for details of the simulations see Appendix D). We then took each trial and computed predictions for the probability of each confidence report using the derived expressions, before randomly drawing a confidence report in accordance with the probability assigned to it. This allowed us to plot confidence simulated from the model, and confidence reports that match the derived predictions.

Fig. 7 shows simulations of confidence using the model (error bars), and confidence based on the derived predictions (error shading). No additional approximations were made in deriving confidence predictions in the interrogation condition. Consistent with this, simulated confidence and the variance of simulated confidence closely match predicted confidence and predicted confidence variance, as functions of response time and unsigned average evidence over the entire-stimulus, both with and without variability in drift-rate scaling.

**Figure 7:**
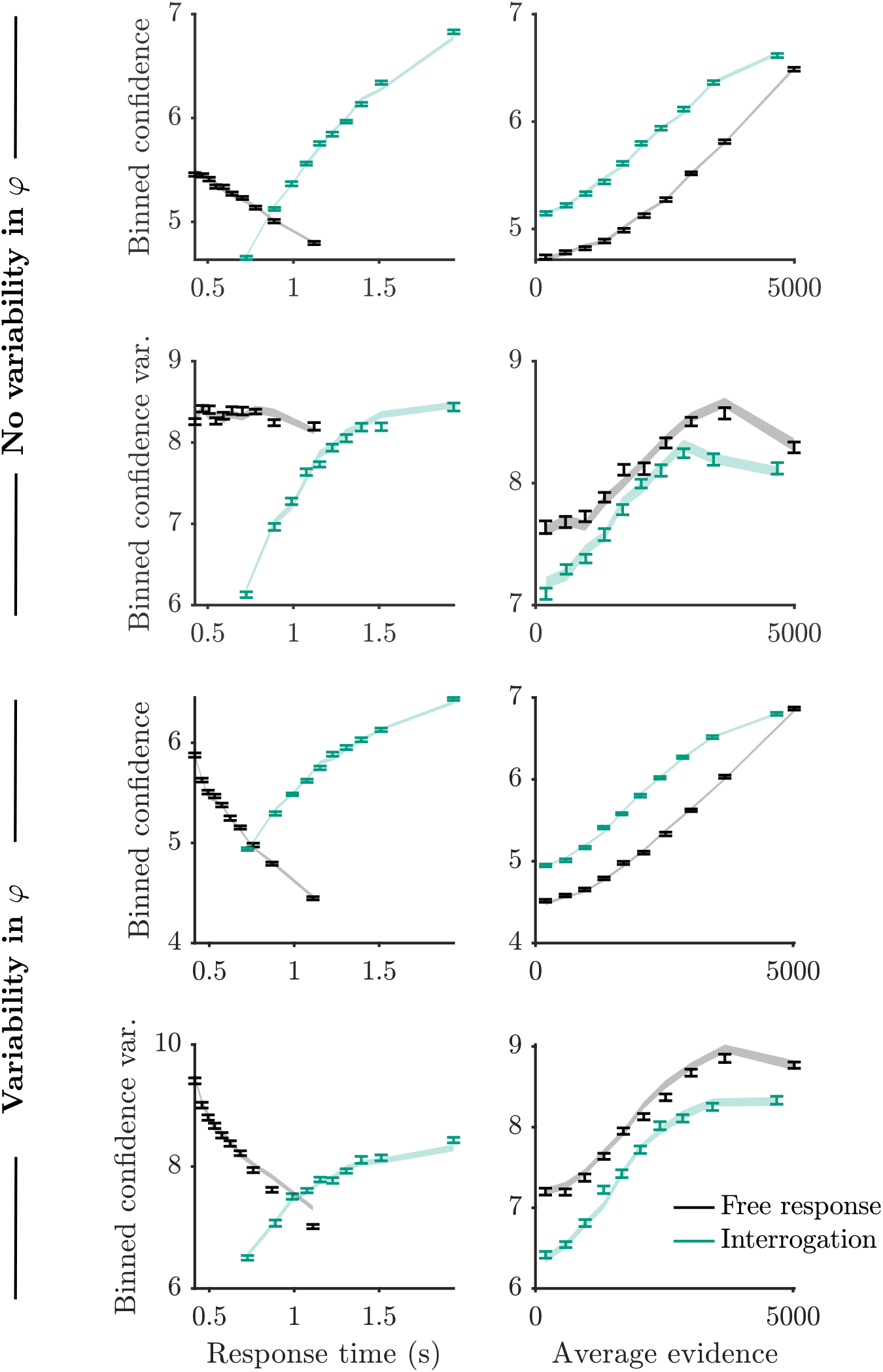
Mean and variance of binned confidence, produced via simulation of the model (error bars), and through the derived predictions (shading). “Average evidence” here refers to the absolute value of the following quantity: The difference in dots summed over all presented frames, divided by the duration of stimulus presentation. Details of the simulation and plotting are provided in Appendix D. Predictions matched the simulation closely. There were some signs that approximations used to derive the predictions lead to a small overestimation of variability in confidence when there is variability in drift-rate scaling (*φ*).

For the free response derivations, we approximated evidence prior to a decision by its average, for the purpose of estimating the drift-rate scaling, and approximated the drift-rate scaling as independent of the average evidence prior to a decision. These approximations are only relevant when the drift-rate scaling is variable. Consistent with this, simulated and predicted confidence closely match in plots corresponding to no variability in drift-rate scaling *φ* (Fig. 7). When variability in drift-rate scaling is present, we can see that the approximations introduce some small discrepancies between simulations and predictions. For example, the predictions appear to overestimate the variability in confidence reports in trials with long response times. For completeness, we note that there were similarly small discrepancies in an observer who used an alternative to Bayesian confidence (Appendix F). Nevertheless, we should always be mindful of the fact that an approximation that works for one model or set of parameters, may not work so well for another model or set of parameters.

## 4 Discussion

Using the normative frameworks of the DDM for decision making and a Bayesian readout for confidence, we derived predictions for the probability distribution over confidence reports, given the response, response time, and stimulus presented on a trial. We considered both the typical case, where response time is under the control of the participant (free response), and the less common case in which the observer has to respond at a particular time (interrogation). In the free response case, where the observer must set decision thresholds to trigger a response, we allow for the use of decision thresholds of arbitrary shape. These results build on the work of Moreno-Bote (2010), by including features that are important in the construction of confidence. Specifically, the derivations account for accumulation of pipeline evidence (Moran et al., 2015), the effect of drift rate variability on pipeline evidence (Pleskac & Busemeyer, 2010), and metacognitive noise (Maniscalco & Lau, 2012, 2016).

Importantly, the derivations cover not only static stimuli but also dynamic stimuli that generate normally distributed fluctuations in evidence signal each frame. The derived expressions only require one evaluation per trial, in contrast to previous approaches that could handle dynamic stimuli but based on evaluation of some function at every time step prior to a decision (e.g. Chang & Cooper, 1970; Smith, 2000; Voss & Voss, 2008; Zylberberg et al., 2018). Reducing computational cost is crucial for making trial-by-trial modelling of dynamic stimuli feasible. Trial-by-trial modelling may provide stronger constraints when fitting models than predictions made for large groups of trials at once (Park et al., 2016), which has until now been the standard approach (see Section 1). Computationally cheap predictions may also allow us to use techniques which require predictions to be evaluated many times, such as cross-validation and Markov chain Monte Carlo (MCMC; Bishop, 2006). A key insight behind our derivations is that it can be much more tractable to model the probability distribution over confidence reports than it is to derive computationally cheap expressions for the decisions (and associated response times) to which the confidence reports relate. This is because in the build up to a confidence report, and specifically after the decision threshold has been crossed, the evolving state of the accumulator follows a normal distribution. In contrast, in the lead up to a decision, the accumulation – constrained to lie at or below a decision boundary – is inherently non-normal.

For readability, we have kept the dependence of the predictions for confidence on model parameters implicit (Section 3), but it will be by adjusting these parameters that we can fit the model described to data (Table 2). Additionally, by constraining parameters to certain values we can construct model variants for model comparison. For example, we could ask whether metacognitive noise is an important source of variability by comparing a model in which we fit all parameters, to a model in which the standard deviation of metacognitive noise is set to zero. There is special flexibility with the decision threshold, because the modeller can choose its shape and how to parameterise it. Using these derivations we have recently compared a variety of models of confidence, including models in which the decision threshold is flat, and models with a decreasing decision threshold (Calder-Travis, Charles, Bogacz & Yeung, 2020).

At the outset we noted that simple expressions may also provide additional insights into the mechanisms responsible for confidence. For example, such expressions may elucidate exactly how we expect different variables to interact to generate confidence, thereby helping us to understand and relate the various patterns that have been observed in confidence data. Consider a situation in which observers report their confidence on a very fine-grained scale in a free response task. In the simple case where the observer scales their readout so that it matches the log-posterior ratio, *x_lp_/K*, the most likely confidence report is (using equation 85) given by the following,

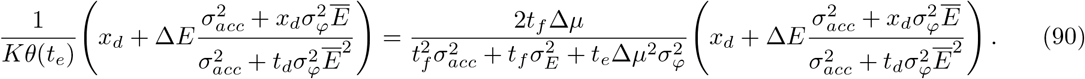

Breaking this expression down, we see that the prefactor multiplying the term in brackets features various sources of variability in the denominator. This factor generates a highly intuitive relationship: When variability increases (and the observer detects this increase in variability) confidence should tend to decrease. Beyond this, the second half of (90), in parentheses, represents a derivation and expression of key principles of the 2DSD model of confidence introduced by Pleskac and Busemeyer (2010). In particular, assuming no variability in drift-rate scaling, the only evidence used for predicting confidence is evidence from the processing pipeline. This is because if drift-rate scaling variability, *σ_φ_*, is zero, the second term within the parentheses,

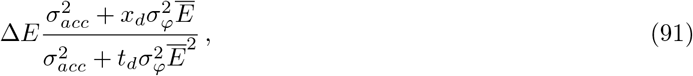

simply reduces to Δ*E*. It seems counterintuitive that the evidence on which the decision was based adds nothing to our prediction for confidence. This occurs because, at the time of the decision, we know that the state of the accumulator is at the decision boundary corresponding to the response made (Fig. 2; Pleskac and Busemeyer, 2010; Yu et al., 2015). Given knowledge of the state of the accumulator at the time of the decision, we do not need to know the evidence presented up to this point.

Drift rate variability adds nuance to this relationship, via the fraction multiplying Δ*E* in (91). This term provides a mathematical description of the fact that, if strong evidence has been gathered by the time of the decision, relative to the time spent deliberating, the drift-rate scaling is likely to be high (Moreno-Bote, 2010), and pipeline evidence will have a big impact on decision confidence (Pleskac & Busemeyer, 2010). On the other hand, if at the time of decision, little evidence has been gathered relative to the time spent deliberating, evidence is accumulating slowly, suggesting a low drift-rate scaling. In turn, this suggests that pipeline evidence will be processed poorly and will have a small effect on confidence. This is why the fraction contains decision time in the denominator, reducing the effect of pipeline evidence, and the height of the threshold at decision time is in the numerator, increasing the effect of pipeline evidence. We are not the first to describe this effect of drift rate variability; it was a central idea in the model for confidence proposed by Pleskac and Busemeyer (2010). Our contribution is to derive an expression for this effect. Moreover, the expression in (90) goes beyond the 2DSD model to include effects of time on confidence stemming from the Bayesian confidence readout used (Moran, 2015; Moreno-Bote, 2010), that are present even in the absence of continued evidence accumulation following a decision, i.e., when Δ*E* is zero. Even in this case, the total time spent processing the stimulus, *t_e_*, still appears in the denominator of the prefactor in (90), and will therefore reduce confidence, consistent with previous findings (Kiani et al., 2014; Murdock & Dufty, 1972).

Beyond formalising our ideas about the relationship between confidence and other variables, a further insight from our expressions for confidence is the integration of empirical findings that previously appeared difficult to explain. In particular, there are inconsistent findings regarding the relationship between confidence and signal strength on error trials. Sanders et al. (2016) found confidence on error trials decreased as signal strength increased, which has been taken as a distinguishing feature of confidence in some studies (Kepecs & Mainen, 2012). However, Kiani et al. (2014) found that confidence on error trials increased with signal strength. In this latter study, participants simultaneously reported their decisions and confidence, and Kiani et al. (2014) suggested that this design choice may have been key to the pattern they observed (see also Desender et al., 2020; Khalvati, Kiani and Rao, 2020). Our derivations support this suggestion. To see why, we look again at our expression for most likely confidence, (90), but now consider the situation in which Δ*E* = 0 (as would be the case if choice and confidence are reported simultaneously). We have,

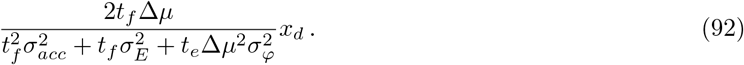

In the case of a flat decision threshold (and, hence, constant *x_d_*), the only variable that changes between trials is *t_e_*, the duration of evidence presentation, which is the same as response time in the free response task. Noting that this term appears in the denominator of the expression, that greater signal strength will lead to faster responses, and that at each level of signal strength the response times for correct and error trials will on average be identical (in the absence of drift-rate variability; Ratcliff and McKoon, 2008; Ratcliff and Rouder, 1998; Shadlen, Hanks, Churchland, Kiani and Yang, 2006), we therefore predict that the most likely confidence report will also be higher on both correct and error trials, due to the lower *t_e_*, when signal strength is greater. A different prediction follows when the observer has time to process pipeline evidence. The processing pipeline contains a considerable amount of information from the stimulus (approximately 400ms of the stimulus prior to response; Ratcliff and McKoon, 2008; Resulaj et al., 2009). On error trials, this evidence will tend to favour the alternative (correct) option, decreasing confidence, and this effect will be stronger when signal strength is high. The implication is that confidence will decrease as signal strength increases.

To test the intuitions gained from studying the equations, we simulated response times, decisions, and confidence reports under two different free response conditions. In one condition, we simulated the task designed by Kiani et al. (2014), setting pipeline evidence to zero. In the other condition, we simulated a processing pipeline containing (a conservative) 100ms of the stimulus prior to response (see Appendix D for details of the simulations). Confidence in errors increased with signal strength when decisions and confidence reports were simultaneous, while confidence in errors decreased with signal strength when observers received pipeline evidence (Fig. 8; statistics in Appendix G).

**Figure 8:**
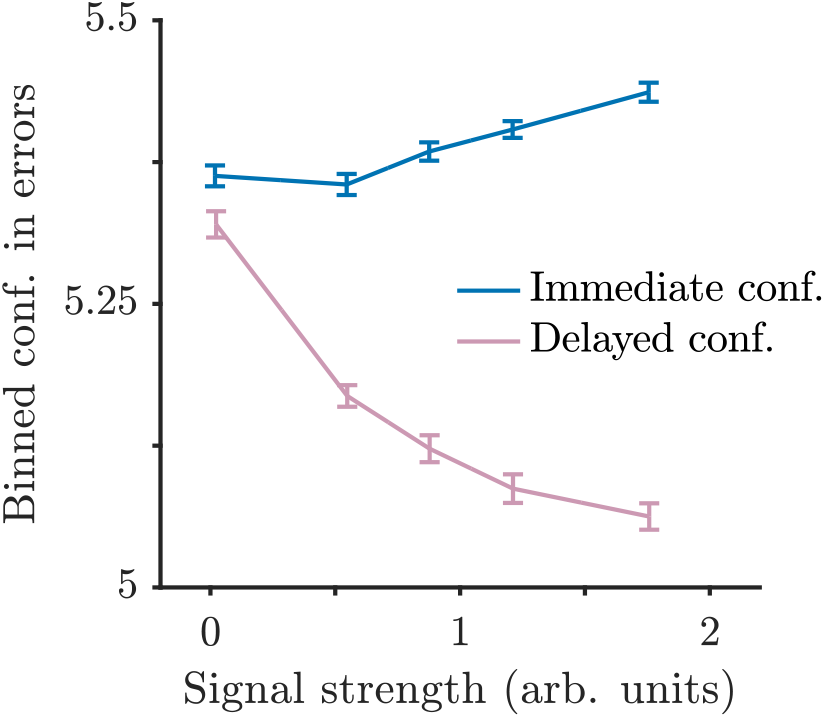
Confidence on free response trials that led to errors, as a function of signal strength. When confidence is reported immediately, and therefore does not reflect evidence measurements in the processing pipeline, confidence on error trials increases with signal strength. On the other hand, when confidence reports are made after a decision, and include 100ms of evidence measurements from the processing pipeline, confidence on error trials decreases with signal strength. Variability in signal strength was generated by using a non-zero value for drift-rate variability. Simulation and plotting details in Appendix D.

In this way, the expressions we derive can provide new insights as well as formalise key intuitions about confidence in tractable expressions for use in future modelling efforts. We should, however, note some important limitations to our approach. First, regarding the scope of the derivations, we have been concerned only with the DDM framework and with a specific class of stimuli. We focus on the DDM as a particularly interesting case due to the normative properties of the diffusion mechanism (see Section 1). Nevertheless, a family of alternatives to the DDM have been studied that assume not just one accumulator, but two (Bogacz et al., 2006; Moreno-Bote, 2010), which may be more or less anti-correlated with each other. It may be possible to use recently derived expressions for the state of the second accumulator at decision time, to extend the approach developed here (Shan et al., 2019), but we leave this for future research. Likewise, we leave for future research whether it is possible to extend the derivations to more general dynamic stimuli, rather than just those with normally distributed evidence fluctuations. In principle, any stimulus for which we can find the optimal observer’s decision and confidence rules could be modelled using the approach we have described. One of the most common stimuli, the random dot motion stimulus, contains dots that move randomly over the course of a trial, creating random fluctuations in evidence for the prevailing motion direction (Kiani et al., 2008; Pilly & Seitz, 2009). If it was possible to characterise the nature of these fluctuations, and derive the optimal observer’s decision and confidence rule, a very large quantity of data could be analysed on a trial-by-trial basis using the approach set out here.

A related limitation is that we have only considered a specific kind of confidence report – a Bayesian readout of the probability of being correct – and have assumed that the observer holds approximately true beliefs about the generative process responsible for their evidence measurements. It is important to note that observers can behave as if they hold a true generative model, and match the behaviour of a Bayesian observer, simply by learning through association the mapping from accumulator state and time to probability correct (Kiani & Shadlen, 2009; Ma & Jazayeri, 2014). Therefore, human confidence reports may be well described by the equations derived above, even if humans are not actually performing Bayesian computations. In addition, the derived equations could be adapted to cover cases where the generative model held by the observer does not perfectly match the true generative model (as in Calder-Travis et al., 2020 for example).

A second set of limitations concern the applicability of our derivations to modelling. In particular, we only make predictions for confidence, not for response times and decisions, and we have not yet accounted for lapses. As detailed above, our choice to derive expressions for confidence alone was a deliberate one, made in order to avoid the difficulties in deriving computationally cheap expressions for responses and response times (see Section 1). Additionally, when fitting confidence reports, we will still model the decision mechanism, in the sense that we will generate estimates for the parameters of this mechanism. Using these parameters we will be able to make predictions for decisions and response times. This will allow us to examine whether a model that fits well to confidence nevertheless generates implausible response and response time data, and will provide an additional check of the model and its assumptions. Regarding lapses, it would be tricky to directly model lapses that affect the response produced (Adler & Ma, 2018; Ratcliff & Tuerlinckx, 2002), because we do not have expressions for the probability distribution over decisions and response times generated by the non-lapse diffusion process. (For similar reasons, it would be difficult to incorporate the idea of variability in the start point of the accumulator, and variability in the duration of the pipeline; Ratcliff and McKoon, 2008; Ratcliff and Tuerlinckx, 2002.) However, it would be straightforward to include a lapse rate parameter that describes some probability that a random confidence report is given.

Notwithstanding the limitations discussed, we believe these derivations will prove useful for the two purposes described at the outset: Supporting deeper insight into the confidence of an important class of observers, and supporting trial-by-trial modelling. We have seen in this discussion that the derived expressions offer the potential to directly explore and understand the relationship between confidence and other variables. This level of explanatory power may be difficult to gain even after running simulations using a wide range of parameter values. The expressions found only require evaluating once per trial, making trial-by-trial modelling of dynamic stimuli more feasible. As a consequence, we hope these results will support efforts to develop models which make ever more precise and sophisticated predictions for behaviour.

## Code availability

All code written for the study will be made publicly available upon publication, and will be accessible through doi:10.17605/OSF.IO/TK3VP.

## Acknowledgements

JCT is grateful for financial support from the Grand Union ESRC Doctoral Training Partnership, and St John’s College, Oxford. This work was supported by Medical Research Council Grant MC UU 00003/1.

## A Product of normal distributions

We will repeatedly encounter a situation where we are interested in the probability distribution over *z* in the following equation,

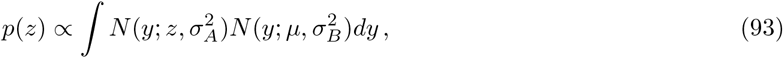

where *N* denotes the normal distribution. The product of two normal distributions is a scaled normal distribution (over *y*; Bromiley, 2014). The scaling is given by,

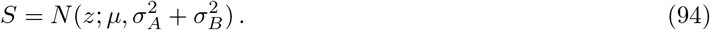

Once we have integrated over *y*, only the scaling is left, as the normal distribution over *y* integrates to one. Hence,

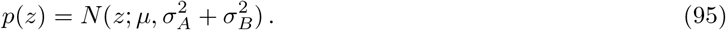

## B Bayesian observer,s policy

At some time *t*, the observer has gathered *N* sensory samples, *δx*_1_, *δx*_2_, *…, δx_N_*, collectively denoted ***δx***. The samples are generated according to (14), (15), and (16). Using Bayes rule the observer can infer the probability of the two possible values for *S*. Using a result from Drugowitsch et al. (2012), Moran (2015) derived the posterior probability over the two options *S* = 1 and *S* = 2, for exactly the case we have. For completeness, we rederive the posterior here.

Given all the evidence measurements so far,

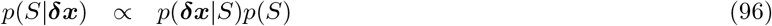

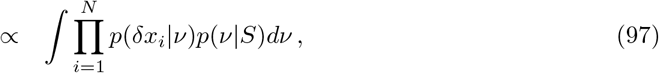

where we have used the flat prior over *S* in (1). Looking at the product in (97), using (16), and using standard formulae for the product of normal distributions (e.g. see Bromiley, 2014) we have,

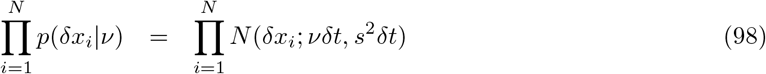

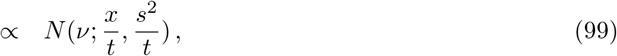

where the proportionality sign indicates proportionality with respect to *ν*, 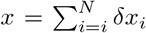, and we have used that 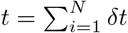. Hence in (97) we have,

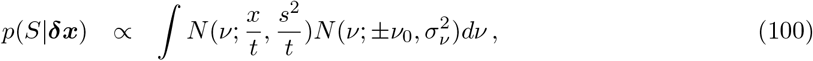

where the sign on *ν*_0_ is determined by *S*. Using the result in Appendix A,

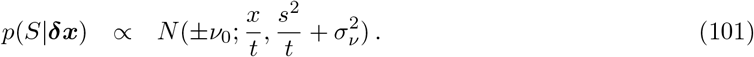

Rearranging, we find the (not scaled) log-posterior ratio is given by,

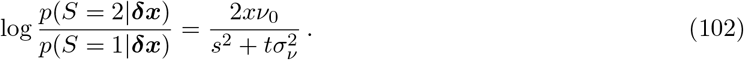

## C Effect of posterior ratio scaling

In the main text we considered a model in which the observer uses a noisy readout of the scaled log-posterior ratio to determine their confidence. A noisy readout of the scaled log-posterior ratio (*x_lp_*), is equivalent to a scaled version of a noisy readout of the unscaled log-posterior ratio (*x_lp_/K*). First consider the readout in the main text in (21):

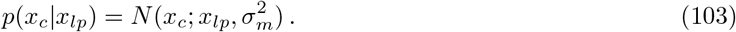

Then consider a noisy readout of the same form but of the unscaled log-posterior ratio. Denote this new readout 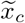. We have,

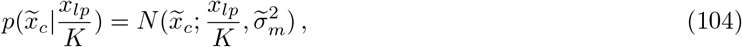

where 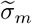 is the standard deviation of the metacognitive noise affecting this readout.

Our claim is that the distribution over 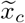 when scaled by K, is identical to the distribution over *x_c_*. Consider a scaled version of 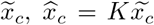. Using the rule for changing the variable of a probability density function, where the relationship between the variables being swapped is monotonic, then,

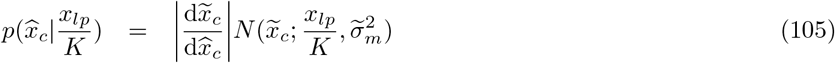

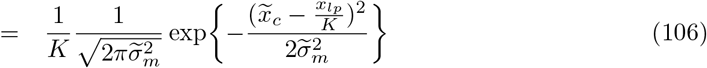

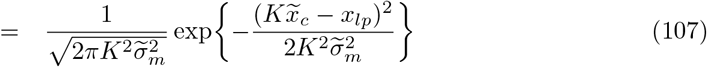

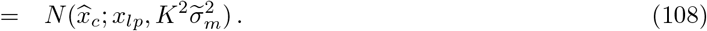

Hence, the scaled readout of the unscaled log-posterior ratio, 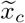, has the same distribution as the readout of the scaled log-posterior ratio, *x_c_*. The metacognitive noise parameters in the two cases are related to each other through 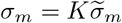.

## D Simulation and plotting details

We simulated the task described in the main text with the parameters shown in Table 3. The only difference between the simulation and the setup described in the main text is that evidence in a frame could not go below 0 or above 3096. Evidence within a frame was resampled until it met these constraints. The midpoint between the means of the two evidence signals, referred to as “reference value” and denoted 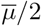, was sampled each trial. It was sampled from a normal distribution centred on 1000, with standard deviation of 100, that was truncated at 500 and 1500.

**Table 3:** Parameters for the task used in the simulation.

| Variable | Expression | Simulation value |
| --- | --- | --- |
| Duration of a frame | $t_f$ | 50ms |
| Max evidence in frame |  | 3096 |
| Min evidence in frame |  | 0 |
| Reference value | $p(\bar{\mu}/2)$ | See text |
| Mean difference between evidence signals | $\Delta\mu$ | 90 |
| Standard deviation of evidence in a frame | $\sigma_E$ | 311 |

The parameters used for the observer are shown in Table 4. Within a simulation, all participants were simulated using the same parameters. Except for Fig. 8, half of the trials were simulated for the free response condition, and half for the interrogation condition. For Fig. 8 all trials were from the free response condition, but two different versions of the free response condition were used, as described in the main text.

**Table 4:** Parameters of the simulated observers.

| Variable | Expression | Fig. 3 | Fig. 7 | Fig. 8 |
| --- | --- | --- | --- | --- |
| Participants per simulation |  | 40 | 40 | 40 |
| Trials per participant |  | 12800 | 12800 | 64000 |
| Pipeline duration | $I$ | 350 ms | 350 ms | 100ms |
| Standard deviation of accumulator noise | $\sigma_{acc}$ | 2860 | 2860 | 2860 |
| Standard deviation of drift-rate scaling | $\sigma_\varphi$ | 1 or 0 (separate simulations) | 1 or 0 (separate simulations) | 0.6 |
| Standard deviation of metacognitive noise | $\sigma_m$ | 0 | 1500 | 1500 |
| Threshold height |  | 2000 | 2000 | 2000 |
| Threshold slope | | 0 | $-800 \text{ s}^{-1}$ | 0 |

A linear decision threshold, symmetric for both options, was used in the free response condition. At the onset of evidence accumulation the threshold was at the value given in Table 4 for threshold height, and it subsequently changed at the rate given by threshold slope. In the interrogation condition, stimulus presentation duration was drawn from a normal distribution of mean 0.8 s and standard deviation 0.3 s, truncated at 0.2 and 4 s.

The evolution of the observer’s accumulator was simulated using small time steps of duration 0.1 ms. Each accumulator increment was determined by sampling from the distribution in (5). Simulation of the accumulation continued until a decision threshold was crossed, and all remaining evidence in the pipeline had been processed. The only exception was the simulation for Fig. 8. For the “Immediate conf.” condition in this plot, accumulation terminated as soon as the decision threshold was reached, consistent with the idea that the manipulation used by Kiani et al. (2014) ensured confidence was not based on pipeline evidence.

Confidence reports were produced in accordance with the model (see (20), (21) and Fig. 2). Where a plot refers to binned confidence, the confidence values produced by the model have been divided into 10 bins of equal numbers of data points on a participant-by-participant basis.

To make predictions for confidence using the derived expressions, we set the values of the parameters in these expressions (*σ_acc_*, *σ_φ_*, *σ_m_*, *I*, threshold height and threshold slope) to the true values used to simulate the data. We set the bin edge parameters (*d_i_*) to the values that were used for dividing the simulated confidence reports into 10 bins.

Prior to plotting, data for the x-axis was binned separately for each participant and data series. For each participant, data series, and bin, the average value of the x-variable was computed. Using these averages, the average value across participants was computed, and this determined the x-location of each bin. 10 bins were used apart from in Fig. 8 where 5 bins were used. The y-variable was calculated separately for each participant and bin, before averaging across participants. Error bars, and the width of error shading, represent *±*1 standard error of the mean across participants.

## E Rearranging interrogation condition result

For the probability distribution over *C* in the interrogation condition, we found in (54) that

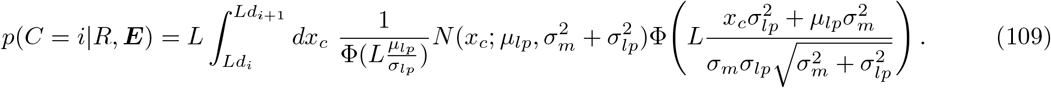

It is possible to rearrange this expression into a form which is easier to evaluate numerically.

Consider a change of variables,

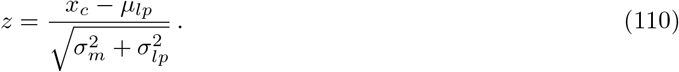

Then we have,

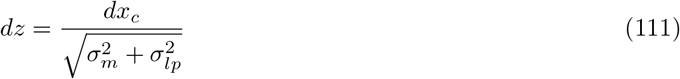

and,

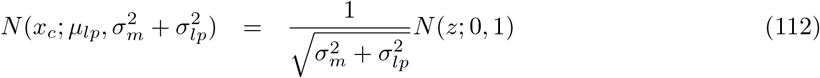

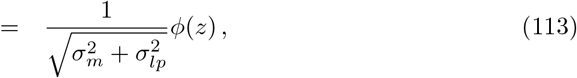

where *ϕ*() indicates the standard normal distribution. Also denote,

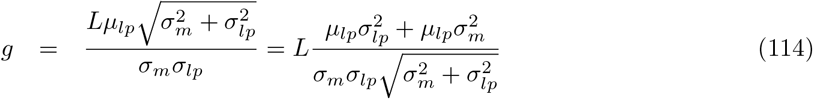

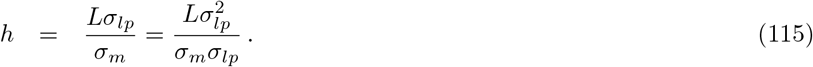

Then,

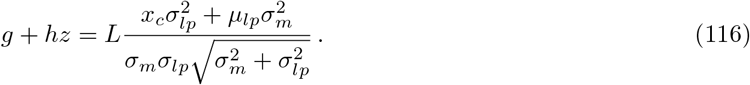

Hence, in (109) we have,

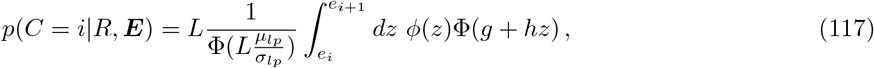

where,

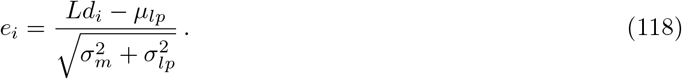

Using result 10010.1 from Owen (1980), gives us,

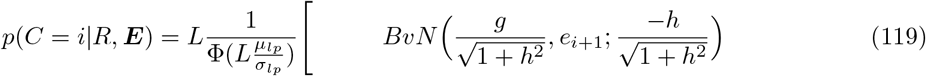

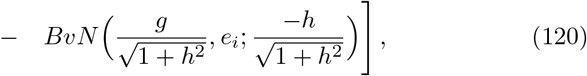

where,

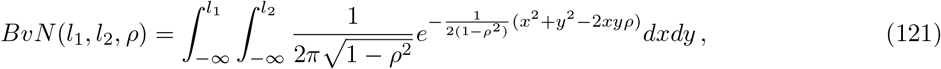

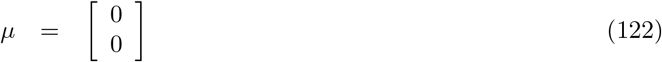

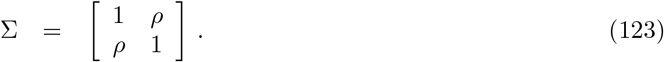

This integral can then be numerically evaluated using standard functions.

## F Testing the approximations in other models

In Fig. 7 we compared the mean and variance of confidence produced via simulations, and confidence produced through the predictions derived in the main text. The aim was to test whether the approximations made during the derivations were reasonable. We found little discrepancy between the simulated and predicted confidence. For completeness, we ran a similar analysis using a model not considered in the main text. Specifically, we considered an observer who did not use a Bayesian readout for confidence. Instead their confidence was directly determined by the final state of the accumulator: More accumulated evidence favouring the choice generated higher confidence. We adjusted the reported derivations to cover this case by setting *θ*(*t*) = 1 (see equation 17).

For this model we found similar discrepancies between the simulations and the derived predictions. Figure 9 provides an example. Again, discrepancies were in general small. Nevertheless, we should bear in mind that just because an approximation works well for some model, or at some parameter values, does not mean it is guaranteed to work well in all models or at all parameter values.

**Figure 9:**
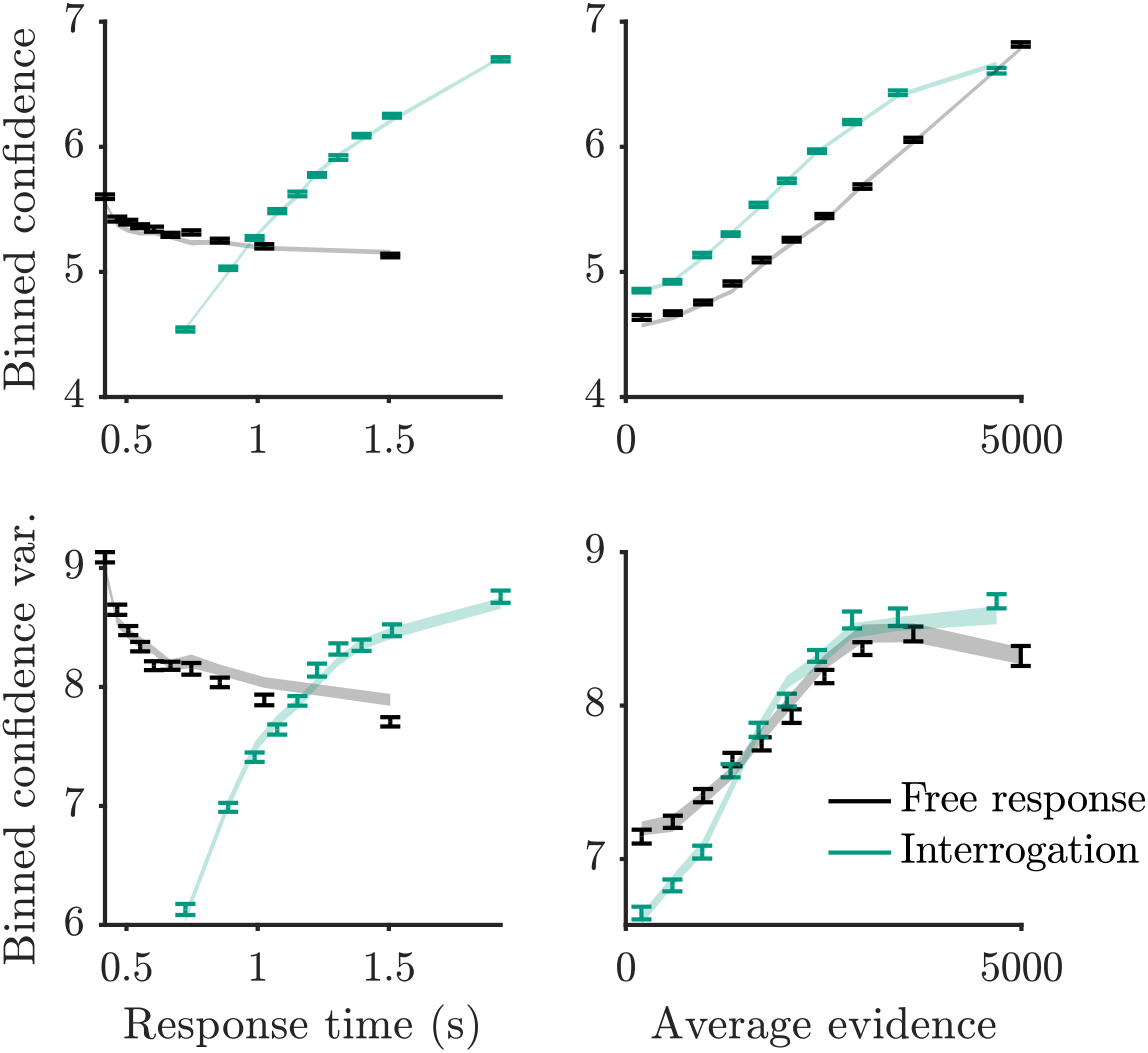
Mean and variance of binned confidence, produced via simulation of an alternative model (error bars), and produced through the derived predictions adjusted for an alternative model (shading). The alternative model featured confidence that was a direct readout of the evidence accumulated, rather than a Bayesian readout for confidence. “Average evidence” here refers to the absolute value of the following quantity: The difference in dots summed over all presented frames, divided by the duration of stimulus presentation.

## G Statistics for Fig. 8

We verified that the patterns observed in Fig. 8 were statistically significant. To do this, for each participant and condition, we calculated the correlation coefficient between the signal strength and binned confidence on error trials. We then compared the correlation coefficients to zero across participants. In the “Immediate confidence” condition binned confidence increased with signal strength (*t*(39) = 7.1, *p* = 1.4 × 10^−8^), while in the “Delayed confidence” condition binned confidence decreased with signal strength (*t*(39) = −17, *p* = 7.7 × 10^−20^).

## Notes

### Competing Interest Statement

The authors have declared no competing interest.

### Summary of Updates

Small improvements throughout.

